# Eukaryotic Initiation Factor 5B (eIF5B)-Driven Translational Control Impacts Oral Squamous Cell Carcinoma Pathophysiology

**DOI:** 10.64898/2026.01.30.702967

**Authors:** Pavan Lakshmi Narasimha, Jinay Patel, Ayan Chanda, Veda Hegde, Beruwalage Hirushika Fernando, Hannah Stephenson, Rebecca Mubaya, Aadarsh Shrestha, Steven C. Nakoneshny, Bo Young Ahn, T. Wayne Matthews, Shamir Chandarana, Robert Hart, Joseph C. Dort, Martin Hyrcza, Emilija Todorovic, Seyed Mehdi Jafarnejad, Pinaki Bose, Nehal Thakor

## Abstract

The non-canonical translation of specific mRNAs has been implicated in oncogenesis and cancer progression. We previously identified eukaryotic Initiation Factor 5B (eIF5B) as a key factor in Internal Ribosome Entry Site (IRES)-mediated translation of a subset of mRNAs encoding anti-apoptotic proteins. Here, we demonstrate that *EIF5B* is predominantly expressed in cancer cells compared to other cell types in the Oral Squamous Cell Carcinoma (OSCC) microenvironment. Higher *EIF5B* mRNA and protein expression are associated with poor patient outcomes. We show that eIF5B depletion in OSCC cells blunted pro-growth, pro-inflammatory, and pro-angiogenic signaling pathways and significantly increased TNF-related apoptosis-inducing ligand (TRAIL)-induced cell death. This is achieved through decreased translation of mRNAs encoding critical factors associated with OSCC pathophysiology. Importantly, the level of interaction of eIF5B with tRNAi^Met^ was significantly higher in OSCC cells compared to non-cancerous fibroblasts. This suggests that OSCC cells (but not non-cancerous fibroblasts) rely heavily on eIF5B for translation initiation. In an in vivo flank xenograft model using nude mice, eIF5B knockdown in UMSCC-29 cells led to a significant reduction in tumor volume compared to control tumors. Also, the immunohistochemical analysis of the xenografted tumor sections demonstrated decreased staining intensity of critical factors associated with OSCC pathophysiology in eIF5B-depleted tumors relative to controls. Collectively, our data demonstrate that OSCC cells are uniquely dependent on eIF5B–tRNAᵢᴹᵉᵗ interactions to sustain translation of pro-survival mRNAs. Targeting eIF5B disrupts these oncogenic programs, sensitizing OSCC cells to apoptosis and suppressing pro-angiogenic and pro-growth signaling.

## Introduction

Head and neck cancers are the sixth most prevalent malignancy worldwide, with over 890,000 new cases and 450,000 deaths reported in 2023 ^1,2^. Among these, oral squamous cell carcinoma (OSCC) is the most common, representing over 90% of cancers arising in the oral cavity, and is notably prevalent with 30,000 new cases in North America ^3,4^. The primary risk factors for OSCC include tobacco and alcohol use, which increase the risk by up to 30 times ^5,6^. Despite the advancements in surgical, chemotherapy and radiotherapy methods, only ∼50% of patients survive beyond 5 years ^7^. These treatment approaches are associated with significant morbidity and have yielded only a 5% improvement in overall survival over the past two decades ^8,9^. This highlights a critical need for novel targeted therapies and prognostic markers based on biological insights.

Dysregulated mRNA translation has been implicated in cancer pathophysiology ^10–12^. Non-canonical translation initiation allows the expression of distinct factors involved in the survival, proliferation, invasion and migration of the cancer cells and tumor angiogenesis ^13,14^. One of the primary mechanisms of non-canonical translation initiation is the internal ribosome entry site (IRES)-mediated translation initiation. Many IRES-containing mRNAs encode oncogenes, growth factors, pro-inflammatory and pro-angiogenic factors, and critical pro-survival/anti-apoptotic proteins ^15^. Thus, IRES-mediated translation initiation provides critical survival and proliferative advantages to the cell under pathophysiological stress conditions.

We previously identified eukaryotic initiation factor 5B (eIF5B), a homologue of bacterial initiation factor 2 (IF2), as a critical factor that drives the non-canonical translation ^16^. In normal cap-dependent translation initiation, eIF5B helps recruit the 60S ribosomal subunit to the 48S pre-initiation complex to form a translation-competent ribosome. However, eIF5B can also deliver and stabilize Met-tRNAi^Met^ into the ribosomal P site during IRES-mediated translation initiation. ^5,6^ As such, eIF5B forms an alternative ternary complex (eIF5B-GTP-Met-tRNAi^Met^) ^17,18^. Under pathophysiological and physiological stress conditions, eIF5B thus takes on a role usually played by the eIF2α–GTP–Met-tRNAi^Met^ complex. Under stress conditions, phosphorylation of eIF2α inhibits its ability to deliver the Met-tRNAi^Met^ to the ribosome, thereby attenuating global mRNA translation. However, a subset of mRNAs evades translation attenuation by switching to IRES-mediated translation initiation. ^7–10^ eIF5B is upregulated in several malignancies and has been implicated in cancer pathophysiology ^11–14^, at least partly by regulating the translation of IRES-containing mRNAs encoding anti-apoptotic proteins ^19^.

In the present study, we interrogated the role of eIF5B in OSCC. We demonstrate that *EIF5B* mRNA is predominantly expressed in tumor cells in the OSCC microenvironment and eIF5B protein is overexpressed in OSCC tumors compared to adjacent normal tissue. We show that high expression of both *EIF5B* mRNA and eIF5B protein are associated with poor outcomes for OSCC patients. Depletion of eIF5B impacts several tumorigenic hallmarks of OSCC cells. By performing polysome profiling experiments, we demonstrate that the translation of mRNAs encoding critical factors affecting OSCC biology was significantly reduced by eIF5B depletion. The level of interaction of eIF5B with initiator tRNA (Met-tRNAi^Met^) is significantly higher in OSCC cells compared to non-cancerous fibroblasts under basal growth conditions. These findings reveal that eIF5B functions distinctly in non-cancerous vs. OSCC cells, with OSCC cells exhibiting a dependency on eIF5B for their survival and proliferation. This highlights eIF5B as a promising therapeutic target in OSCC.

## Materials and Methods

### Patient Cohorts

Genomic and clinical data were acquired for multiple OSCC cohorts. The Cancer Genome Atlas (TCGA) provided transcriptomic profiles for 275 treatment-naive tumors and 26 normal mucosal samples. Raw/normalized expression values and clinical information were obtained from UCSC Xena ^20^. In addition, *EIF5B* mRNA and eIF5B protein expression levels were assessed using Clinical Proteomic Tumor Analysis Consortium (CPTAC) HPV-negative OSCC patient cohort data, consisting of 46 treatment-naïve tumors and 22 normal mucosal samples ^21^. A single-cell RNA-seq dataset was obtained from Puram *et. al.* (GSE103322, ^22^). The dataset, comprising ∼6000 cells from 18 HPV-negative OSCC patients, was analyzed using the Seurat package (version 5.0.1) ^23^. After excluding samples from metastatic sites, a total of 4486 cells were retained, including 1706 malignant and 2780 non-malignant cells. Quality control steps included filtering cells with low gene counts and normalizing the data to mitigate technical variations. Following clustering and cell-type annotation, specific gene expression levels were evaluated across different cell populations within the TME.

Our institutional cohort comprised 175 surgically resected, treatment-naive OSCC cases diagnosed between 2009 and 2013 and accrued within the Ohlson Research Initiative (ORI) at the University of Calgary, with a median follow-up of 5.8 years (current to July 2024). Cohort characteristics included a median age of 62.5 years and prospective outcome tracking (Table 1). Our study adheres to REMARK guidelines ^24^ for tumor biomarker validation and received ethical approval from Alberta’s Health Research Ethics Board (HREBA-Cancer Committee #HREBA.CC-16-0113). All procedures complied with Canada’s Tri-Council Policy Statement for human subject research.

**Table 1:** Demographics of OSCC patients in Ohlson TMA cohort. P-values were calculated using the Wilcoxon rank-sum test for continuous variables and Fisher’s exact test for categorical variables (two-sided).

| **Variable**<br>n (%) unless<br>otherwise<br>indicated | **All**<br>**Patients** | **EIF5B**<br>**Low** | **EIF5B**<br>**High** | **p-**<br>**value** |
| --- | --- | --- | --- | --- |
| Number | 176 | 51 | 125 | - |
| Sex |  |  |  | 0.864 |
| Female | 65 (36.9) | 18 (27.7) | 47 (72.3) |  |
| Male | 111 (63.1) | 33 (29.7) | 78 (71.3) |  |
| Median age at<br>diagnosis, years<br>(IQR) | 64.1 (21.4) | 58.2 (18.5) | 65.0 (21.0) | **0.046** |
| Pathologic<br>T stage |  |  |  | 0.131 |
| T1/T2 | 105 (59.6) | 35 (33.3) | 70 (66.7) |  |
| T3/T4 | 63 (35.8) | 15 (23.8) | 48 (76.2) |  |
| Missing | 8 (4.6) | 1 (12.5) | 7 (87.5) |  |
| Pathologic<br>N stage |  |  |  | 0.804 |
| Node-negative | 66 (37.5) | 17 (25.8) | 49 (74.2) |  |
| Node-positive | 78 (44.3) | 23 (29.5) | 55 (71.5) |  |
| Missing | 32 (18.2) | 11 (34.4) | 21 (65.6) |  |
| TNM Stage |  |  |  | 1 |
| Stage I/II | 43 (24.4) | 12 (27.9) | 31 (72.1) |  |
| Stage III/IV | 101 (57.4) | 28 (27.7) | 73 (72.3) |  |
| Missing | 32 (18.2) | 11 (34.4) | 21 (65.6) |  |
| Extracapsular spread |  |  |  | 0.314 |
| Present | 51 (29) | 18 (35.3) | 33 (64.7) |  |
| Absent | 125 (71) | 33 (26.4) | 92 (73.6) |  |
| Differentiation |  |  |  | 0.0932 |
| Well | 38 (21.6) | 6 (15.8) | 32 (84.2) |  |
| Moderate | 100 (56.8) | 32 (32) | 68 (68) |  |
| Poor | 35 (19.9) | 13 (37.1) | 22 (62.9) |  |
| Missing | 3 (1.7) | 0 (0) | 3 (100) |  |
| Lymphovascular Invasion |  |  |  | 0.402 |
| Absent | 135 (76.7) | 37 (27.4) | 98 (72.6) |  |
| Present | 20 (11.4) | 6 (30) | 14 (70) |  |
| Missing | 21 (11.9) | 8 (38.1) | 13 (61.9) |  |
| Perineural Invasion |  |  |  | 0.366 |
| Absent | 99 (56.2) | 33 (33.3) | 66 (66.7) |  |
| Present | 54 (30.7) | 14 (25.9) | 40 (74.1) |  |
| Missing | 23 (13.1) | 4 (17.4) | 19 (82.6) |  |
| Alcohol consumption history |  |  |  | 0.522 |
| Never drinker | 32 (18.2) | 11 (34.4) | 21 (65.6) |  |
| Ever drinker | 134 (75.1) | 38 (28.4) | 96 (71.6) |  |
| Missing | 10 (5.7) | 2 (20) | 8 (80) |  |
| Smoking history |  |  |  | 0.461 |
| Never smoker | 48 (27.3) | 16 (33.3) | 32 (66.7) |  |
| Ever smoker | 127 (72.2) | 35 (27.6) | 92 (72.4) |  |
| Missing | 1 (0.5) | 0 (0) | 1 (100) |  |
| Median pack-years smoked (IQR) | 20 (35) | 17 (35) | 20 (35) | 0.403 |

### GTEx data integration and cross-tissue comparison

Gene expression for EIF5B was extracted from three sources: (i) the TCGA HPV-negative OSCC tumor cohort, (ii) matched normal tissues from the same OSCC study, and (iii) non-diseased reference tissues from the Genotype-Tissue Expression (GTEx) project. All expression values were already on a comparable scale, log2(TPM+1). For GTEx, per-sample EIF5B expression values were merged with GTEx sample metadata using the GTEx sample identifier, and samples with missing or non-informative tissue annotations (e.g., “not provided”) were excluded. GTEx tissues were then grouped by the metadata field defining histological tissue type (“Subsite”). For visualization, OSCC tumor samples were labeled “OSCC” and matched non-tumor samples were labelled “Matched_normal”. The plotting order was pre-specified to display OSCC and Matched_normal first, followed by all GTEx tissues sorted alphabetically. Expression distributions were summarized using boxplots per Subsite, with overlaid jittered points to show per-sample values. To evaluate whether EIF5B expression in OSCC tumors differed from each reference tissue distribution, pairwise comparisons were performed between the OSCC group and each non-OSCC Subsite (Matched_normal and each GTEx tissue) using a two-sided Wilcoxon rank-sum test. Multiple testing across these comparisons was controlled using the Benjamini–Hochberg (BH) false discovery rate adjustment. Statistical significance was displayed on the plot as asterisk annotations above each Subsite box only when BH-adjusted p-values met conventional thresholds (*, <0.05; **, <0.01; ***, <0.001).

### Statistical Analysis

Baseline clinicopathologic characteristics were compared between EIF5B expression groups using the Wilcoxon rank-sum test for continuous variables and Fisher’s exact test for categorical variables (two-sided). Where categorical contingency tables met chi-square assumptions (all expected cell counts ≥5), Pearson’s chi-square test was used; otherwise, Fisher’s exact test was applied. Continuous variables are reported as median (IQR) unless otherwise specified. Overall survival (OS), disease-specific survival (DSS), and progression-free interval (PFI) were analyzed using Kaplan–Meier estimates with two-sided log-rank tests. For Kaplan–Meier curves with continuous variables, a cut-point determined by the method outlined by Contal and O’Quigley was utilized.^25^ Univariable and multivariable Cox proportional hazards models were used to estimate hazard ratios (HRs) with 95% confidence intervals (CI). For analyses involving multiple hypothesis tests, p-values were adjusted using the Benjamini–Hochberg (BH) procedure to control the false discovery rate (FDR). Unless otherwise stated, BH-adjusted p-values (q-values) < 0.05 were considered statistically significant. Missing data were handled using complete-case analysis for each model (i.e., individuals with missing values for a given variable were excluded only from analyses involving that variable); no imputation was performed. The protein expression status of University of Calgary OSCC patients was dichotomized based on low (0/1) and high (2/3) eIF5B protein levels for Kaplan–Meier curves. Spearman’s ρ was calculated for the correlation between *EIF5B* mRNA and protein expression. *EIF5B* mRNA expression was compared between tumour and matched normal tissue using the Wilcoxon signed-rank test. An unpaired t-test was performed for the *in vitro* assays. All statistical analyses were performed using GraphPad Prism (GraphPad, California) or R version 4.4.3. Unless specified, all quantified data show the mean ± standard error of the mean (SEM) for 3 biological replicates. An unpaired, two-tailed t-test determined statistical significance without assuming equal variance. The significance level was set at a p-value of 0.05.

### Immunohistochemistry (IHC)

Tissue slides were deparaffinized, rehydrated, and subjected to antigen retrieval using citrate buffer. Endogenous peroxidase activity was quenched with a peroxidase block, and slides were blocked and permeabilized using rodent block with 0.2% Triton X. Slides were incubated for one hour with anti-eIF5B (Proteintech 13527-1-AP) followed by an HRP-conjugated goat anti-rabbit secondary antibody (Dako EnVison+ kit, K4065), and finally for 2 min with 3,3′-diaminobenzidine tetrahydrochloride (DAB) to visualize bound antibodies. Slides were counterstained with hematoxylin, dehydrated, and mounted. Multiple antibody concentrations for eIF5B (1:100, 1:200, 1:500 and 1:1000) were used to optimize staining for eIF5B protein in control mouse muscularis and normal oral cavity squamous epithelium core samples (Fig. S1). To determine the ideal antibody concentration that would enable the scoring of differential protein expression levels within TMA cores, the study pathologist (ET) evaluated stained slides. After staining optimization, the final primary antibody concentration of 1:200 for eIF5B was selected. Images of representative cores were captured using the EP50 camera attached to an IX50 inverted microscope (Olympus, Canada). The pathologist (DI) assigned a score to each TMA; a score of 0 denoted nonexistent staining, while scores of 1, 2, or 3 indicated increasing intensity of positive staining for eIF5B protein. A pathologist scored the staining on a scale of 0 to 3, and 1 was considered low expression, while 2 and 3 were high expression. By calculating the patient’s maximum score, discrepancies between cores from the same case were settled ^26^.

### Cell Culture and Reagents

CAL-33 and BJ-5ta (immortalized fibroblast) cell lines were obtained from American Type Culture Collection (ATCC; Virginia, USA), and UMSCC-1 and UMSCC-29 were obtained from the University of Michigan (Michigan, USA). All lines were propagated in Dulbecco’s high-modified Eagle’s medium (DMEM, HyClone) with 4 mM L-glutamine, 4,500 mg/L glucose, and 1 mM sodium pyruvate, supplemented with 10% heat-inactivated fetal bovine serum (Gibco), and 1% penicillin-streptomycin (Gibco). Cells were incubated at 37°C in an incubator supplemented with 5% CO_2_. Reverse transfections were performed using Lipofectamine RNAiMAX (Invitrogen) according to the manufacturer’s instructions and scrambled non-specific negative control siRNA (siC; Qiagen) and siGenome SMARTpool of siRNA targeting eIF5B (M-013331-01-0010), eIF2A (M-014766-01-0005) & eIF2D (M-003680-01-0005) were obtained from Dharmacon.

### Cell Viability Assay

Cells were seeded into 96-well plates at the following densities: CAL-33 (1000 cells/well), UMSCC-29 (3000 cells/well), UMSCC-1 (5000 cells/well) and BJ-5ta (5,000 cells/well). Cells were reverse-transfected with either control siRNA (siC) or eIF5B (si5B), and after 24 hours, they were treated with vehicle (DMEM media) or TRAIL (100 ng/mL) for an additional 72 hours. Cell viability was assessed using the Alamar Blue assay (Resazurin sodium salt; Sigma-Aldrich) according to the manufacturer’s protocol, and fluorescence (excitation 560 nm, emission 590 nm) was measured using a Cytation 5 plate imager (BioTek); values were normalized to vehicle-treated controls.

### Flow cytometry

CAL-33/UMSCC-29/UMSCC-1 and BJ-5ta cells were seeded at 200-300,000 cells/well and reverse-transfected in six-well dishes with siC or si5B. After 72 h, TRAIL or a vehicle was added. After a further 4 h, cells were harvested by trypsinization, followed by the Apoptosis assay according to the manufacturer’s protocol (STEMCELL Technologies #100-0338). Briefly, cells were resuspended in PBS containing 2% FBS after trypsin treatment. Then cells were centrifuged and resuspended in Annexin V buffer containing 2% FBS. Finally, cells were suspended in Annexin V buffer at 1×10^6^ /mL concentration and 100 µl was removed, followed by the addition of 5 µL of FITC and/or 7AAD before analyzing using Flow cytometry (BD FACSAria III).

### Apoptotic Assay

OSCC cells were seeded at 1,000 cells/well and reverse-transfected in a 96-well plate. 100 ng/mL TRAIL was added after 92 h of incubation. 4 h after TRAIL addition, cells were rinsed in 1x PBS, followed by adding 1x Annexin binding buffer with 1 μg/mL Hoechst and Annexin V-FITC. Cells were imaged at 20 X magnification using the Cytation 5 plate imager. Cells were imaged using a DAPI or GFP filter to visualize Hoechst and Annexin-V.

### Western Blotting

Cells were seeded at 100,000 - 300,000 cells/well in a 6-well plate and reverse-transfected using control siRNA (siC; Qiagen) or siRNA specific for eIF5B (si5B; Dharmacon) for 72-96 hours. 100ng/mL TRAIL was added 92 hours after transfection and harvested after 4 more hours. Cells were harvested in radioimmunoprecipitation assay (RIPA) lysis buffer with protease inhibitors. Equal amounts of proteins (typically 20 μg; 30 μg for cFLIP_s_ and cleaved caspase 7) were resolved by SDS-PAGE and transferred onto a nitrocellulose membrane (GE Healthcare). Nitrocellulose membranes were blocked with milk powder dissolved in 0.1% phosphate buffer saline (typically 5% wt/vol; 0.5% wt/vol for cFLIP_s_ and cleaved caspase 7). Individual proteins were incubated with the primary antibodies listed in Supplementary Table 1 overnight with appropriate dilutions, followed by incubation with appropriate HRP-conjugated secondary antibodies. ECL reagent was then added to the blots for visualization and the immunoblots were imaged in an AI600 image (GE Healthcare) and densitometry was performed using the AI600 analysis software. For the 4EGI-1 experiment, BJ-5ta cells were seeded at 300,000 cells/well in a 6-well plate, and siRNA transfection was performed as described above using si-C or si-eIF5B. 48h after transfection, cells were treated with 25 µM 4EGI-1 or a vehicle control (DMSO). 24 h post 4EGI-1 treatment, cells were lysed using RIPA. Western blot analysis was performed for eIF5B, XIAP, and cIAP1, as mentioned above.

### RNA-IP

100 μl of protein G Dynabeads™ (Invitrogen) was incubated with eIF5B-specific antibody (1:50 in 0.02 % 1X PBST) for 4 h at 4 °C. After 4 h of incubation, the beads were washed with PBS. 10-cm culture plates were seeded with BJ-5ta/UMSCC-29. Upon reaching 80% confluency, the plates were washed twice with ice-cold 1X PBS, and the cells were lysed using 1 mL RIPA buffer containing protease, phosphatase, and RNAse inhibitors. The cell lysates were incubated with antibody-coated Dynabeads^TM^ on a rotary shaker at 4°C. The beads were then separated using a magnetic rack and washed thrice with 500 μl of cold nuclease-free PBS. RNA was extracted from the beads using the phenol:chloroform method. RT-qPCR was performed as described by Ho JJD., et al. ^27^ to quantify eIF5B-bound tRNA_i_^Met^. The tRNA_i_^Met^ levels were normalized to input. The qPCR primer sequences are mentioned in Supplementary Table S1.

### BrdU Incorporation Assay

CAL-33 (1,000 cells/well), UMSCC-29 (3,000 cells/well), UMSCC-1 (5,000 cells/well), and BJ-5ta (5,000 cells/well) cells were seeded and reverse-transfected with siRNA in 96-well plates. After 24-48 hours, 1x BrdU was added according to the manufacturer’s instructions (2750, Sigma-Millipore). BrdU was incorporated for 24 hours. Following incubation, cells were fixed, and DNA was denatured using the provided fixing solution. Incorporated BrdU was detected using an anti-BrdU antibody and a peroxidase-conjugated secondary antibody, with color development achieved using TMB substrate. Absorbance was measured at 450 nm and 550 nm using a Cytation 5 plate imager (BioTek), and results were interpreted as proportional to cell proliferation.

### Collagen Invasion Assays

24 hours before beginning the assay, serum-containing media was removed from OSCC cells and replaced with low serum media. OSCC cells were then seeded at 200,000-250,000 cells/well and reverse-transfected with either siC or si5B into collagen inserts in a 24-well plate in serum-free media with serum-containing media on the other side of the insert. Cells were incubated for 48 h followed by microscopy images and extraction was performed according to the manufacturer’s instructions (QCM Collagen Cell Invasion Assay, 24-well 8μm, calorimetric; ECM551 from Sigma Aldrich). Briefly, the inserts were dipped in water (MilliQ) and the inside of the inserts was cleaned using the cotton tips provided in the kit. Microscopy images were taken using Cytation 5, inserts were stained with Cell stain (Crystal violet), incubated for 10 min, and OD was read at 560 nm.

### In vitro scratch assay

UMSCC-29, CAL-33, and UMSCC-1 cells (5 × 10⁵ per well) were seeded in 12-well plates, treated with control siC or si5B for 48 h, and grown to near confluency in complete growth medium. Cells were serum-starved overnight in 0.2% FBS medium to stop replication without impacting cell survival. A 200 μL pipette tip was used to create a linear scratch in the monolayer, followed by PBS washing and incubation in 0.2% FBS medium (30 h for all cell lines) at 37°C under 5% CO₂. Scratch closure was monitored using an Olympus CKX53 and/or Cytation 5 microscope (10x objective) with 0 h and 30 h post-incubation. Five images per well were captured along the scratch axis, and scratch widths were measured at three positions per image (15 total/condition) using ImageJ (v1.53, NIH). Data represent mean ± SEM from three independent experiments, with cell line sources, reagent details, and imaging parameters fully documented for reproducibility.

### Angiogenic Biomarker Analysis

OSCC and BJ-5ta cells were seeded (100,000 cells/well) and reverse-transfected using siC or si5B in 6-well plates. After 72 hours, the media was replaced with serum-free media. TRAIL (100 ng/mL) was added 4 h before harvesting. Spent media was collected, centrifuged, and analyzed using the Human Angiogenesis & Growth Factor 17-Plex Discovery Assay® (Eve Technologies). This assay simultaneously measured 17 angiogenesis and growth factor-related markers, including Angiopoietin-2, BMP-9, EGF, Endoglin, Endothelin-1, FGF-1, FGF-2, Follistatin, G-CSF, HB-EGF, HGF, IL-8, Leptin, PLGF, VEGF-A, VEGF-C, and VEGF-D, with sensitivities ranging from 0.2 – 42.8 pg/mL.

### Endothelial Tube Formation Assays

Human umbilical vein endothelial cells (HUVEC) were seeded at 50,000 cells/well in a 96-well plate coated with Matrigel. Cells were supplemented with spent OSCC media collected for the angiogenic biomarker assay. Branch formation was allowed to take place for 6 - 8 h. Brightfield microscopy images were taken at 6 - 8 h. Analysis of tube formation characteristics was performed using ImageJ, using the available angiogenesis macro ^28^.

### Polysome Profiling and RT-qPCR

UMSCC-29 and CAL-33 cells were seeded at 1.5–2.0 × 10⁶ cells per 10-cm dish and transduced with either control shRNA (shC) or shRNA targeting eIF5B (sh-eIF5B) using five plates per condition. After 24 h, the medium was replaced with puromycin-containing medium (2 µg/mL) to select transduced cells, and selection was continued for an additional 48 h. The resulting control and eIF5B-depleted populations were pooled and lysed on ice in RNA/polysome lysis buffer (0.3 M NaCl, 15 mM MgCl₂·6H₂O, 15 mM Tris-HCl pH 7.4, 1% Triton X-100, 0.2% sodium deoxycholate if desired, 2 µL SUPERase-In, and 5.5 µL of 1% cycloheximide per 1.1 mL buffer) to preserve ribosome–mRNA complexes. Equal amounts of clarified lysate were layered onto 10–50% linear sucrose gradients; an aliquot of each lysate was reserved for total RNA isolation and for verification of eIF5B knockdown by immunoblotting. Gradients were centrifuged in an SW41 rotor at 39,000 rpm for 90 min at 4 °C, and 1-mL fractions were collected using a density-gradient fractionation system while continuously monitoring absorbance to generate polysome profiles. RNA from each fraction was extracted with acid phenol:chloroform (5:1, Ambion), ethanol-precipitated, and resuspended in RNase-free water ^19^. Equal volumes of RNA from each fraction were reverse-transcribed using the qScript cDNA synthesis kit (Quanta Biosciences) ^19^, and RT-qPCR was performed on a CFX-96 real-time thermocycler (Bio-Rad) with LunaScript RT SuperMix (New England Biolabs) and gene-specific primers are mentioned in Supplementary Table S1. The percentage mRNA distribution was quantified as previously described ^29^.

### In vivo xenograft mouse model

All animal procedures were performed in accordance with the guidelines of the Canadian Council on Animal Care and were approved by the Institutional Animal Care Committee at the University of Lethbridge (Protocol #2507). UMSCC-29 cells transduced with either control (shC) or eIF5B-targeting (sh5B) shRNA were injected subcutaneously into athymic nude mice (Crl:NU(NCr)-Foxn1nu; male, 5–6 weeks of age; 1 × 10^6 cells per mouse). Tumor growth was monitored and caliper measurements were obtained at least three times per week. Tumor volumes were analyzed using a linear mixed-effects model [Volume = (Length X Width^2^)/2], and differences were considered statistically significant at P ≤ 0.05. Mice were euthanized at predefined humane endpoints, and tumors were excised, processed, and sectioned for downstream analyses. Mouse tumors were removed, fixed overnight in Neutral Buffered formalin (NBF), embedded in paraffin and sectioned as shown. 5-µm tissue sections were stained with H&E for histological analysis. Brightfield images were acquired using a NanoZoomer at 40X. Formalin-fixed paraffin-embedded tumor sections were deparaffinized in xylene, rehydrated through graded ethanol, and rinsed in water, followed by heat-mediated antigen retrieval using citrate buffer (10mM, pH 6 for 10min) containing Tween-20 using a pressure cooker. After cooling, sections were encircled with a hydrophobic barrier, washed in Tris-buffered saline with Tween-20, and incubated with a peroxidase block and a protein blocking solution supplemented with Triton X-100. Slides were then incubated with anti-eIF5B antibody (1:200 in antibody diluent), for 1 h at 4°C, washed, rest of the antibodies were probed overnight XIAP (1:50), Bcl-xL (1:100), VEGFA (1:100), HIF1α (1:200) and p-EGFR (1:100) and exposed to a DAKO HRP-conjugated anti-rabbit polymer for 30 min, before visualization with 3,3′-diaminobenzidine (DAB) substrate for approximately 5 min and hematoxylin counterstaining for 1 min. Sections were rinsed in lukewarm running water, dehydrated through ascending ethanol concentrations, cleared in two changes of xylene, mounted with DPX and coverslips, and allowed to cure overnight at room temperature before sealing the edges with nail polish and storage at 4°C.

## Results

### High eIF5B Expression Correlates with Poor Survival Outcomes in OSCC

We first evaluated the prognostic significance of eIF5B in OSCC using publicly available datasets from GTEx, TCGA, and CPTAC. In GTEx dataset EIF5B (transcript) expression was broadly detectable across all evaluated tissues, with a median expression level in OSCC tumors that was comparatively elevated relative to most GTEx tissue groups, whereas matched normal oral mucosa exhibited a slightly lower central tendency than tumors (Fig. 1A). TCGA (Fig. 1B) and CPTAC (Fig. 1C) data analysis revealed that EIF5B mRNA expression was significantly higher in OSCC tumors compared to normal tissues, a pattern that was consistently observed across other head and neck squamous cell carcinoma (HNSCC) subsites (Supple. Fig. 1A & B). . Additionally, analysis of publicly available single-cell RNA-Seq ^23^ data demonstrated that *EIF5B* mRNA is predominantly expressed in malignant epithelial cells relative to other cell types in the OSCC microenvironment (Fig. 1D). Survival analysis using TCGA data revealed that high *EIF5B* mRNA expression was associated with significantly worse overall survival (Fig. 1E). In HNSCC, we observed a similar trend when EIF5B expression was high (Supple. Fig. 1C) Moreover, higher *EIF5B* mRNA also correlated with higher pathologic node (N) status, suggesting a link to metastatic progression in OSCC (Fig. 1F). Also, eIF5B protein levels were significantly elevated in OSCC and HNSCC (Fig. 1G & Supple. Fig 1D; respectively).

**Figure 1:**
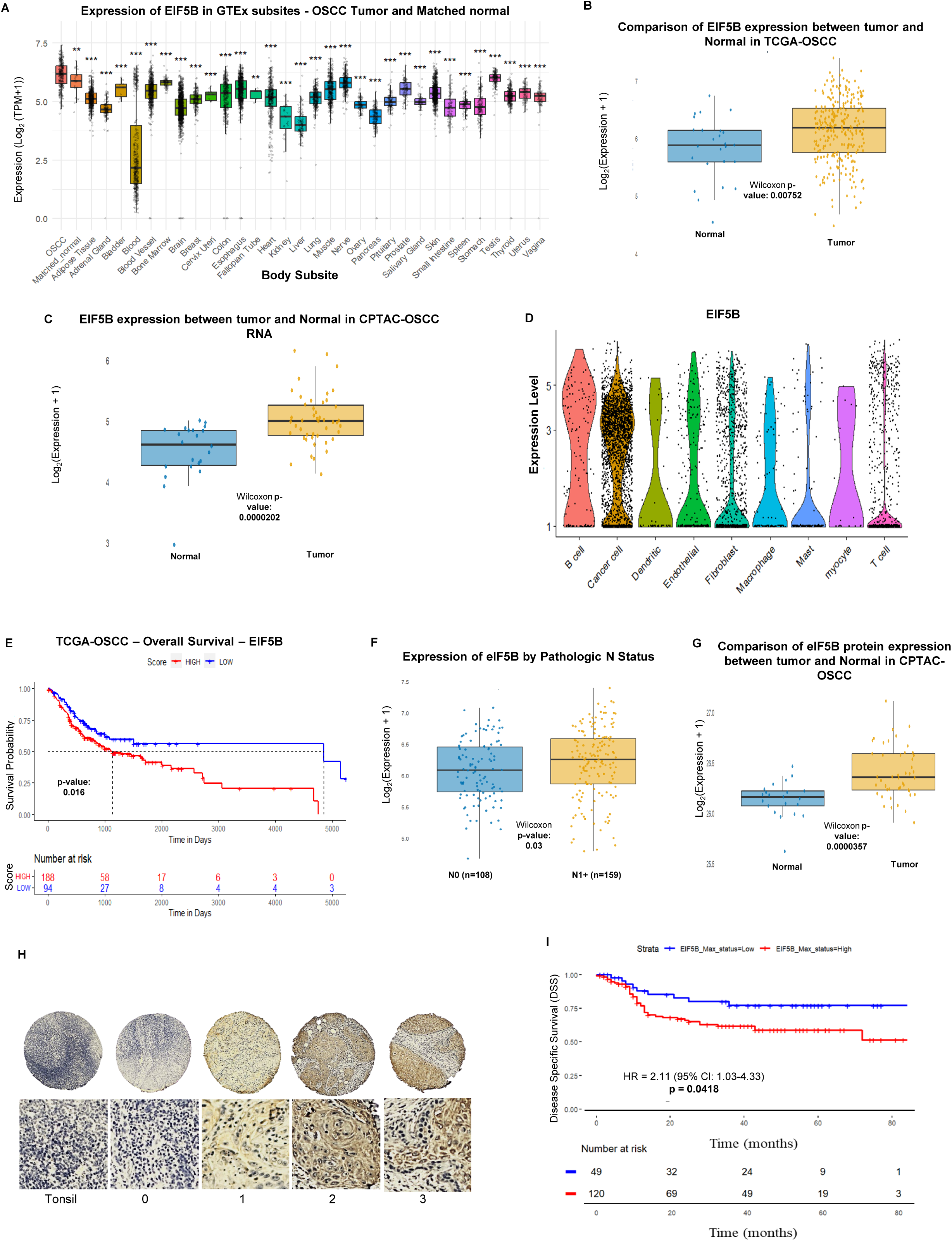
eIF5B expression is associated with poor clinical outcomes of OSCC patients. **(A)** EIF5B expression in OSCC tumors, matched normal tissues, and GTEx reference tissues. Boxplots show the distribution of EIF5B expression (log2[TPM+1]) for OSCC tumors (OSCC), matched non-tumor samples (Matched_normal), and GTEx tissues grouped by histological subsite. Center lines denote medians, boxes represent the interquartile range (IQR), and whiskers extend to 1.5×IQR, points represent individual samples. For each non-OSCC group, expression was compared to OSCC using two-sided Wilcoxon rank-sum tests and p-values were adjusted for multiple comparisons using the Benjamini–Hochberg method. Asterisks denote BH-adjusted significance levels displayed only for significant comparisons (*, <0.05; **, <0.01; ***, <0.001). **(B)** *EIF5B* mRNA expression in OSCC tumors and normal samples. Boxplot showing log2(expression + 1) of *EIF5B* mRNA in normal (blue) and tumor (yellow) samples from TCGA-OSCC (The Cancer Genome Atlas - Oral Squamous Cell Carcinoma) HPV-negative cohort. p=0.00752; Wilcoxon analysis. Each dot represents an individual sample. **(C)** Boxplot showing log2(expression + 1) of *EIF5B* mRNA in normal (blue) and OSCC tumor (yellow) samples from the CPTAC-OSCC cohort. p<0.0002; Wilcoxon analysis. **(D)** Analysis of publicly available single-cell RNA-Seq data from OSCC patients (HPV negative cohort) (Puram SV et al 2017, *Cancer Cell*). **(E)** Kaplan-Meier curve depicting overall survival of TCGA-OSCC patients stratified by EIF5B expression levels. Patients with high EIF5B expression (red curve) display significantly reduced overall survival compared to those with low EIF5B expression (blue curve), as indicated by a log-rank p-value of 0.016. The x-axis represents survival time in days, and the y-axis indicates survival probability. The number of patients at risk at specified time points is provided below the plot for each group. Hazard ratios and p-values quoted are from univariate coxph models; values in square brackets indicate 95% confidence intervals. **(F)** *EIF5B* mRNA expression is significantly elevated in tumors with lymph node metastasis. Box plot comparing *EIF5B* gene expression levels between tumors without lymph node metastasis (N0, n=108) and tumors with pathologic lymph node metastasis (N1+, n=159), analyzed using the Wilcoxon rank-sum test (p = 0.03). **(G)** Analysis of the Clinical Proteomic Tumor Analysis Consortium (CPTAC) data demonstrated that tumor samples (yellow) expressed significantly higher eIF5B (protein) compared to normal samples (blue). p<0.0001; Wilcoxon analysis. **(H)** The OSCC tissue microarrays (TMAs) (176 tumors with 3 cores each) were stained with the eIF5B antibody. Normal tonsil tissue was used as a control. TMAs were scored for the levels of immuno-reactivity based on the staining intensity within the tissue samples as 0: no expression, 1: low expression, 2: high expression, and 3: high expression. Disagreements between cores from the same case were resolved by taking the maximum score for that patient. Scale bars indicate 50 μm. **(I)** Kaplan–Meier survival curves for high (2/3) vs. low (0/1) eIF5B expression reveal that patients with high eIF5B expression had significantly poorer Disease-Specific Survival (DSS). Hazard ratios and p-values quoted are from univariate coxph models, values in brackets indicate 95% confidence intervals.

To further investigate the correlation between eIF5B protein levels and patient outcomes, we analyzed commercially available patient-derived HNSCC TMAs (34 normal and 180 HNSCC tissues; tissuearray.com). We demonstrated that the muscularis of the mouse colon did not stain positively for eIF5B (Supple. Fig. 1E). This was expected because the muscularis does not express high levels of eIF5B protein and served as a negative control. Compared to cancer adjacent tongue (Supple. Fig. 1F) and cancer adjacent larynx tissue (Supple. Fig. 1H), tongue cancer (Supple. Fig. 1H and K), and larynx cancer tissues (Supple. Fig. 1J) expressed high levels of eIF5B protein. Because these commercial TMAs lacked survival data, we performed analysis using TMAs from the ORI at the University of Calgary that contained 176 tumors (each with 3 cores) (Fig. 1H). We analysed the clinicopathological characteristics of OSCC patients from the ORI cohort, stratified by eIF5B expression (Table 1). Demographic analysis revealed significant age differences between eIF5B-low (median 58.2 years) and eIF5B-high (65.0 years) groups (p=0.046), with no sex predilection (p=0.864). Univariate Cox regression identified extracapsular spread (HR=3.045, p<0.001), nodal involvement (HR=2.37, p=0.001), and TNM stage (HR=2.518, p=0.004) as key predictors of poor overall survival. Alcohol/smoking history showed no survival correlation, while perineural invasion trended toward significance for progression-free interval (p=0.076). High eIF5B protein levels were associated with significantly worse disease-specific survival (DSS) in OSCC patients (Figure 1I) and a trend towards worse overall survival (OS) and progression-free interval (Supple. Fig. 2). High eIF5B expression independently predicted worse disease-specific survival in multivariate analysis (HR=2.308, p=0.047) (Table 2). Multivariate models confirmed TNM stage as the strongest predictor of disease-specific mortality (HR=7.027, p=0.015). Collectively, these results indicate that elevated EIF5B transcript and eIF5B protein levels in OSCC are associated with aggressive tumor biology and adverse patient outcomes, independent of traditional risk factors.

**Table 2:** Cox regression for Survival Conditions among OSCC patients in the Ohlson TMA cohort.

|  | Univariate Analysis |  |  |  |  |  |  |  |  |
| --- | --- | --- | --- | --- | --- | --- | --- | --- | --- |
|  | Overall Survival |  |  | Disease Specific Survival |  |  | Progression Free Interval |  |  |
| Variable | HR | CI | p-value | HR | CI | p-value | HR | CI | p-value |
| Sex | 1.074 | (0.692-1.667) | 0.751 | 0.882 | (0.507-1.536) | 0.658 | 1.034 | (0.613-1.744) | 0.901 |
| Age at Diagnosis | 1.041 | (1.024-1.059) | **1.62E-06** | 1.031 | (1.01-1.053) | **0.0036** | 1.017 | (0.998-1.037) | 0.0717 |
| Median Pack Years Smoked | 1.008 | (0.995-1.021) | 0.217 | 1.016 | (1-1.033) | **0.0473** | 1.013 | (0.999-1.028) | 0.0766 |
| Smoking History | 1.34 | (0.819-2.194) | 0.244 | 1.801 | (0.902-3.596) | 0.0956 | 1.81 | (0.941-3.481) | 0.0752 |
| Alcohol History | 1.061 | (0.615-1.831) | 0.832 | 1.326 | (0.621-2.828) | 0.466 | 1.577 | (0.747-3.326) | 0.232 |
| Lymphovascular Invasion | 1.055 | (0.969-1.149) | 0.22 | 1.07 | (0.981-1.168) | 0.127 | 1.086 | (0.995-1.185) | 0.064 |
| Perineural Invasion | 1.528 | (0.95-2.456) | 0.0802 | 1.687 | (0.92-3.093) | 0.0912 | 1.352 | (0.773-2.365) | 0.29 |
| Extracapsular Spread | 3.045 | (1.848-5.017) | **1.24E-05** | 2.869 | (1.496-5.503) | **0.00152** | 2.438 | (1.354-4.389) | **0.00297** |
| Pathologic T Stage | 1.022 | (0.648-1.611) | 0.926 | 1.624 | (0.922-2.861) | 0.093 | 1.304 | (0.79-2.151) | 0.299 |
| Pathologic N Stage | 2.37 | (1.412-3.979) | **0.00109** | 3.847 | (1.833-8.074) | **0.000367** | 2.724 | (1.463-5.071) | **0.00158** |
| TNM Stage | 2.518 | (1.345-4.713) | **0.00389** | 10.518 | (2.537-43.613) | **0.00118** | 3.77 | (1.602-8.87) | **0.00237** |
| EIF5B Expression | 1.513 | (0.918-2.496) | 0.105 | 2.11 | (1.028-4.331) | **0.0418** | 1.703 | (0.923-3.139) | 0.0882 |
|  | Multivariate Analysis |  |  |  |  |  |  |  |  |
|  | Overall Survival |  |  | Disease Specific Survival |  |  | Progression Free Interval |  |  |
| Extracapsular Spread | 2.608 | (1.442-4.716) | **1.51E-03** | 1.829 | (0.887-3.768) | 0.102 | 1.818 | (0.916-3.611) | 0.0875 |
| Pathologic N Stage | 1.374 | (0.636-2.97) | 0.419 | 1.525 | (0.64-3.633) | 0.341 | 1.427 | (0.623-3.27) | 0.4 |
| TNM Stage | 1.598 | (0.672-3.805) | 0.289 | 7.027 | (1.458-33.862) | **0.0151** | 2.604 | (0.903-7.513) | 0.0766 |
| EIF5B Expression | 1.714 | (0.942-3.12) | 0.0777 | 2.308 | (1.013-5.259) | **0.0466** | 2.048 | (0.981-4.275) | 0.0562 |

### The expression of anti-apoptotic proteins and TRAIL-induced cell death in OSCC cells is modulated by eIF5B

Having established that eIF5B overexpression correlates with adverse clinical outcomes in OSCC, we next examined its functional role in tumor cell survival. We hypothesized that eIF5B promotes OSCC cell persistence by supporting the selective translation of anti-apoptotic mRNAs and conferring resistance to TRAIL-mediated apoptosis. eIF5B depletion, in combination with TRAIL treatment, led to a significant reduction in alamarBlue^TM^ activity (viability) in UMSCC-29, CAL-33, and UMSCC-1 cells (Fig. 2A, C, D). To check if eIF5B depletion enhanced TRAIL-mediated apoptotic cell death, we performed flow cytometry with Annexin V-FITC/7-AAD stained OSCC cells. Compared to the control cells (siC), TRAIL treatment (siC + TRAIL) did not robustly enhance apoptosis in UMSCC-29 cells (Fig. 2B). Depletion of eIF5B resulted in robust enhancement in early apoptosis (Fig. 2B). Both early and late apoptosis were further enhanced when eIF5B was depleted in cells treated with TRAIL (Fig. 2B). Likewise, eIF5B depletion greatly enhanced TRAIL-mediated apoptosis in CAL-33 (Supple. Fig. 3A) and UMSCC-1 (Supple. Fig. 4A) cells. This was further confirmed by performing fluorescence microscopy experiments that showed robust Annexin V labelling under eIF5B depletion + TRAIL treatment conditions in UMSCC-29, CAL-33, and UMSCC-1 cells (Supple. Fig. 5).

**Figure 2:**
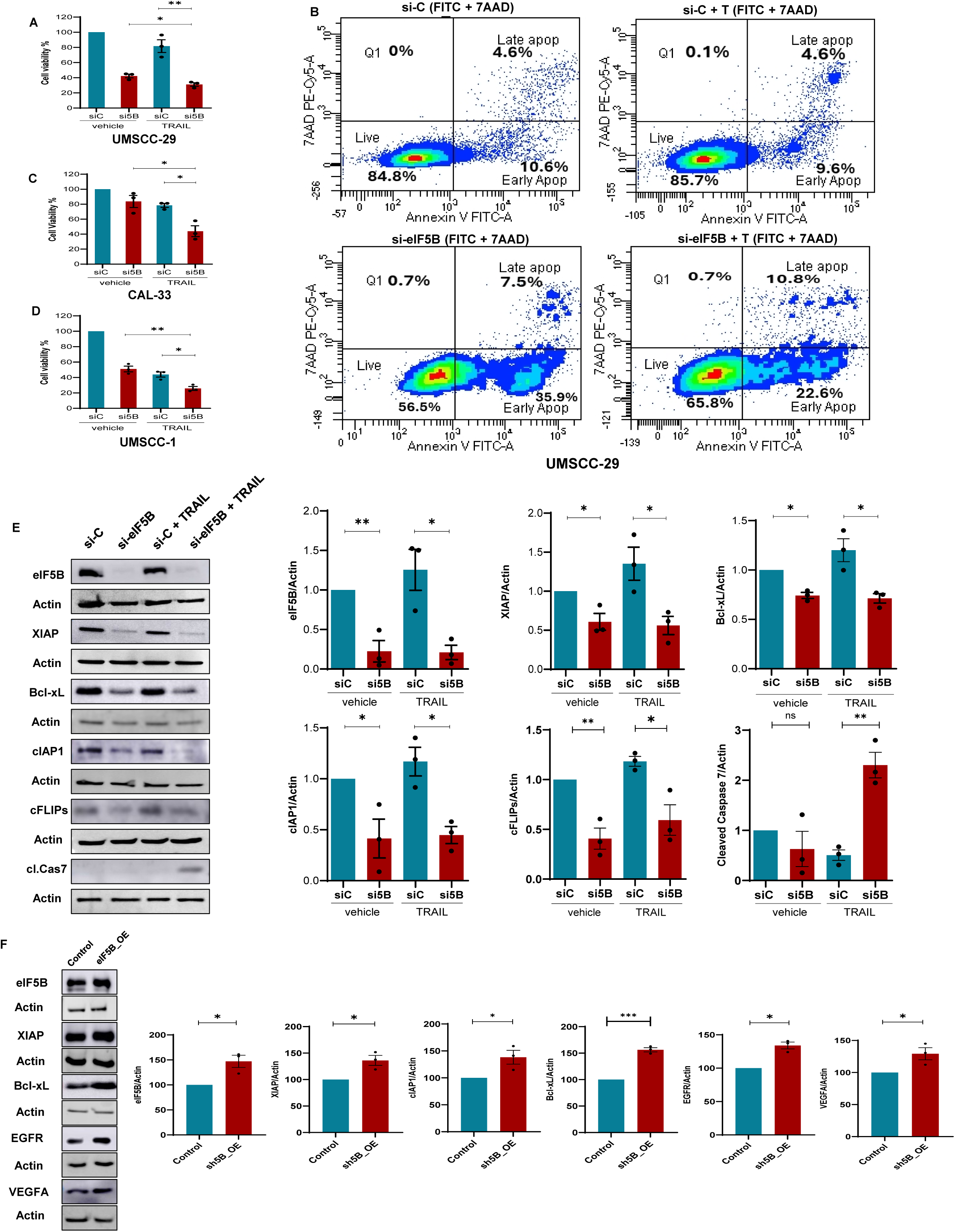
eIF5B regulates apoptosis in UMSCC-29 cells via regulating the expression of distinct anti-apoptotic proteins. **(A)** UMSCC-29, **(C)** CAL-33 and **(D)** UMSCC-1 cells were transfected with either a non-specific control siRNA (siC) or an eIF5B-specific siRNA pool (si-eIF5B). Following 24 h of transfection, cells were treated with either vehicle control or TRAIL (100 ng/mL). Cell viability was assessed after an additional 72 hours using alamarBlue^TM^ assays. **(B)** Annexin V and 7-AAD staining and flow cytometric analysis of transfected cells. The top two panels, siC and siC + TRAIL, the bottom two panels, si5B and si5B + TRAIL, were harvested at 96 h post-transfection and treated with Annexin V + 7AAD. Contour plots are divided into quadrants Live (lower left; cells staining negative for both 7-AAD and annexin V), Q1 (upper left; cells staining positive for 7-AAD but negative for annexin V), Early apoptosis (lower right; staining annexin V positive and 7-AAD negative), and Late apoptosis (upper right; staining positive for both 7-AAD and annexin V) **(E)**. Control or eIF5B-depleted cells were treated with TRAIL and western blot analysis was performed. TRAIL (100 ng/mL) treatment was limited to 4 h before harvesting to prevent excessive cell death of OSCC cells. **(F)**. eIF5B was exogenously overexpressed in UMSCC-29 cells using an eIF5B-expressing plasmid vector, and the levels of XIAP, Bcl-xL, EGFR, and VEGFA were monitored. Data are presented as mean ± SEM from three independent biological replicates using t-test, with statistical significance indicated as follows: *, p < 0.05; **, p < 0.01; ***, p < 0.001.

eIF5B depletion decreased levels of anti-apoptotic proteins (under both control and TRAIL treatment conditions), such as XIAP, Bcl-xL, cIAP1, and c-FLIPs, in UMSCC-29 (Fig. 2E), and CAL-33 cells (Supple. Fig. 3B). In contrast, exogenous overexpression of eIF5B in UMSCC-29 (Fig. 2F) and Cal 33 (Supple. Fig. 3C) significantly enhanced expression of distinct anti-apoptotic proteins. Unlike UMSCC-29 and CAL-33 cells, cIAP1 was not affected by eIF5B depletion in UMSCC-1 cells (Supple. Fig. 4B). As eIF2A is also known to interact with initiator tRNA ^30,31^, we depleted eIF2A from CAL-33 and UMSCC-29. We did not observe any decrease in XIAP, Bcl-xL, cIAP1, and c-FLIPs levels in eIF2A-depleted OSCC cells (Supple. Fig. 6). In all three OSCC cell lines, caspase-7 was not activated until we combined eIF5B depletion with TRAIL treatment (Fig. 2A; Supple. Fig. 3B, Supple. Fig. 4B). In contrast, depletion of eIF5B in BJ-5Tta fibroblasts did not show a decrease in almarBlue^TM^ activity under TRAIL treatment conditions (Fig. 3A). Also, there was no activation of TRAIL-mediated apoptosis in eIF5B-depleted BJ-5ta cells (Fig. 3B). Notably, eIF5B depletion did not alter the levels of anti-apoptotic proteins such as XIAP, Bcl-xL, cIAP1, and c-FLIPs in BJ-5ta cells (Fig. 3C). Furthermore, we did not observe notable activation of caspase 7 in BJ-5ta cells, indicating a lack of apoptotic cascade initiation under eIF5B depletion + TRAIL treatment conditions. We further confirmed that BJ-5ta cells expressed DR4 and DR5 (TRAIL receptors) (Fig. 3C). These findings suggest that BJ-5ta cells possess the necessary receptors for TRAIL binding, yet unlike OSCC cells, eIF5B depletion from BJ-5ta cells had no notable effect on the anti-apoptotic proteins or survival under TRAIL treatment. Interestingly, when we inhibited eIF4E in BJ-5ta using a chemical inhibitor, 4EGI, and combined it with eIF5B depletion, we saw a significant decrease in XIAP levels (Fig. 3D). This suggests that depletion of eIF5B alone is not sufficient to decrease the levels of XIAP in non-cancer BJ-5ta cells. It also suggests that XIAP is likely predominantly expressed using canonical translation initiation in non-cancer cells, in contrast to OSCC cells.

**Figure 3:**
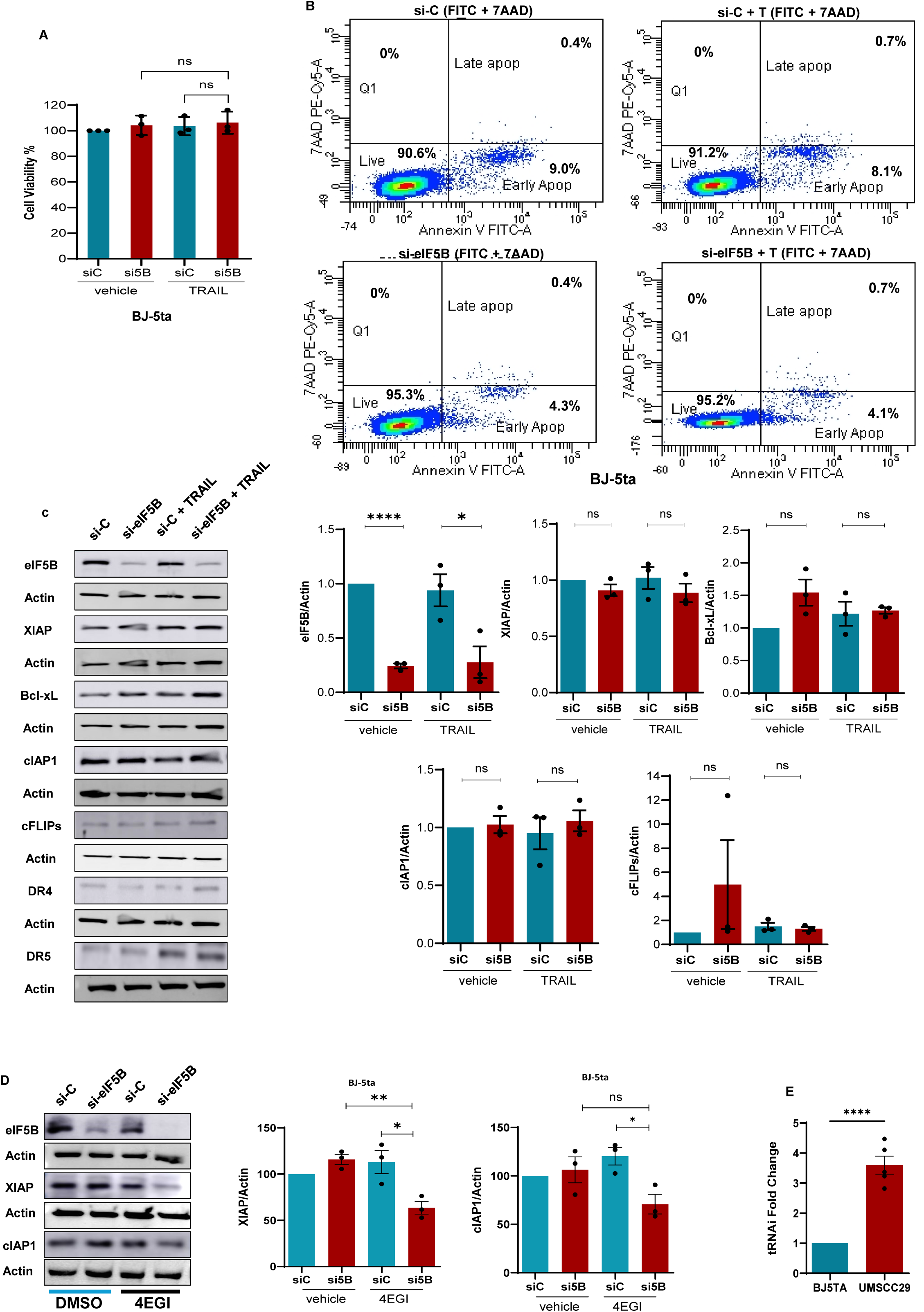
eIF5B depletion does not activate apoptosis in non-cancer immortalised fibroblast cells. BJ-5ta cells were transfected with either a non-specific control siRNA (siC) or an eIF5B-specific siRNA pool (si-eIF5B). **(A)** Following 24 h of transfection, cells were treated with either vehicle control or TRAIL (100 ng/mL). Cell viability was assessed after an additional 72 h using alamarBlue^TM^ assays. **(B)** Annexin V and 7-AAD staining and flow cytometric analysis of transfected cells. The top two panels, siC and siC + TRAIL, the bottom two panels, si5B and si5B + TRAIL, were harvested at 96 h post-transfection and treated with Annexin V + 7AAD. Contour plots are divided into quadrants Live (lower left; cells staining negative for both 7-AAD and annexin V), Q1 (upper left; cells staining positive for 7-AAD but negative for annexin V), Early apoptosis (lower right; staining annexin V positive and 7-AAD negative), and Late apoptosis (upper right; staining positive for both 7-AAD and annexin V). **(C)** Representative immunoblots probing for eIF5B, XIAP, Bcl-xL, cIAP1, cFLIPs, DR 4, DR 5, and β-actin (internal control) are shown. (**D)** Control and eIF5B-depleted BJ-5ta cells were treated with the 4EGI-1 inhibitor to inhibit cap-dependent translation. Levels of XIAP were monitored under control and the 4EGI-1 treatment conditions. **(E)** eIF5B-bound RNA immunoprecipitations (RIPs) followed by qRT-PCR measurements of input-normalized tRNAiMet levels from BJ-5ta vs UMSCC-29. Data are presented as mean ± SEM from three independent biological replicates, with statistical significance indicated as follows: *, p < 0.05; **, p < 0.01; ***, p < 0.001.

To examine why, when compared to BJ-5ta cells, expression of anti-apoptotic proteins in OSCC cells is more prone to eIF5B depletion, we performed RNA-IP experiment for eIF5B and probed for the initiator tRNA. Interestingly, we found that the interaction of eIF5B with the initiator tRNA is several-fold higher in OSCC cells compared to BJ-5ta cells (Fig. 3E). Therefore, we propose that OSCC cells are more dependent on the eIF5B-mediated mechanism of initiator tRNA delivery to the ribosome. This would be particularly important for the translation of the IRES-containing mRNAs such as VEGF, HIF-1α, Bcl-xL, XIAP, cIAP1, and cFLIPs. As a result, depletion of eIF5B robustly decreased the levels of these proteins in OSCC cells compared to BJ-5ta cells.

### The proliferative, migratory, and invasive properties of OSCC cells are controlled by eIF5B

Since eIF5B depletion sensitized OSCC cells to apoptosis and reduced anti-apoptotic protein expression, we investigated whether eIF5B similarly influences other malignant phenotypes. We next examined its role in regulating proliferation, migration, invasion, and angiogenic potential in OSCC cells. We began by performing BrdU incorporation assays in OSCC cells to assess changes in proliferation upon eIF5B depletion. The results demonstrated a significant reduction in BrdU incorporation in eIF5B-depleted UMSCC-29 (Fig. 4A), CAL-33 (Supple. Fig. 7A), and UMSCC-1 (Supple. Fig. 8A) cells. This suggests that the depletion of eIF5B has a significant impact on the proliferation of OSCC cells. To investigate whether this effect was specific to cancerous cells, we conducted similar assays in non-cancerous fibroblast (BJ-5ta) (Fig. 4B) cells. Interestingly, eIF5B depletion in BJ-5ta cells resulted in a significant increase in BrdU incorporation, suggesting that the decrease in proliferation due to eIF5B depletion is specific to OSCC cells. These findings highlight the differential role of eIF5B in regulating proliferation between cancerous and non-cancerous cells.

**Figure 4:**
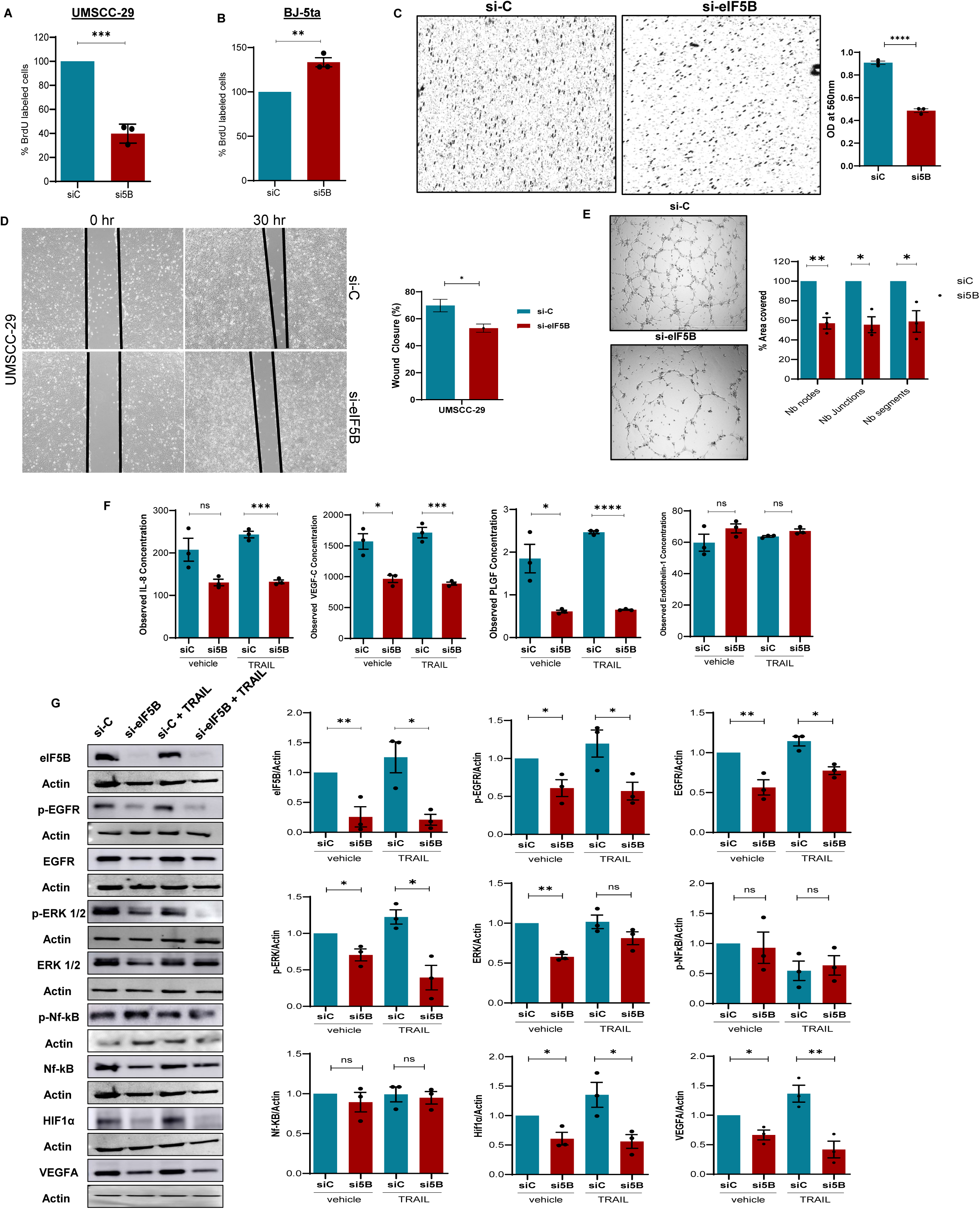
eIF5B depletion decreases the proliferation, invasion, migration and angiogenesis of UMSCC-29 cells. BrdU incorporation assay was used to measure cell proliferation. Quantification shows the percentage of BrdU-labeled cells in control (siC) and si5B-treated groups. Data are presented as mean ± SEM; ***p < 0.001. **(A)** UMSCC-29 **(B)** BJ-5ta **(C)** Collagen-based invasion assay was used to evaluate cell invasiveness in control (siC) and si5B-treated groups. Cells were seeded onto collagen matrices and allowed to invade for a defined period (48-72 hours). The number of invading cells was quantified and expressed as the mean ± SEM. Microscopy images of invading cells **(C; left panel)**, quantification **(C; right panel)**. Phase contrast microscopy images showing wound area in siC/si5B-treated OSCC cells at the time of scratch introduction (T = 0 h) & at T = 30 h wound closure **(c; top panel)**. Quantification of the percentage of wound area covered after 30 hours, normalized to wound area at the time of scratch introduction **(c; lower panel)**. **(E)** Representative images of the endothelial HUVEC cells tube formation assay treated with conditioned media from siC- or si5B-transfected OSCC. ImageJ angiogenesis macro software quantified Junctions, segments, and nodes. OSCC cells were transfected with either control siRNA (siC) or eIF5B-specific siRNA (si5B) and incubated for 96 hours. **(F)** The conditioned media for the control and eIF5B-depleted OSCC cells were analyzed for angiogenic biomarkers **(lower panel)** using EVE Technologies. **(G)** Representative images of immunoblots quantified probing for eIF5B, p-EGFR, EGFR, p-ERK, ERK, p-NF-kB, NF-kB, HIF1α, VEGFA, and β-actin (internal control). Data are presented as mean ± SEM from three independent biological replicates, with statistical significance indicated as follows: *, p < 0.05; **, p < 0.01; ***, p < 0.001, and ****, p< 0.0001.

To examine the effect of eIF5B depletion on the invasive ability of OSCC cells, a collagen invasion assay was performed. The grey-scaled cells in the control condition indicated a substantially higher number of invading cells in control conditions compared to the eIF5B-depleted UMSCC-29 cells and a significant reduction in the invading cells was observed in eIF5B-depleted UMSCC-29 cells compared to control (Fig. 4C). Likewise, the invasion of eIF5B-depleted CAL-33 (Supple. Fig. 7B) and UMSCC-1 (Supple. Fig. 8B) was significantly decreased. These results suggest that eIF5B depletion appears to impair the invasive ability of OSCC cells. Wound healing assays were performed to investigate the effect of eIF5B depletion on the migratory ability of OSCC cells. A scratch was covered nearly 100% by the control OSCC cells, indicating robust cell migration. In contrast, eIF5B depletion in UMSCC-29 cells led to a significant reduction in wound closure by approximately 40% (Fig. 4D). We also observed a similar trend in CAL-33 (Supple. Fig. 7C) and UMSCC-1 (Supple. Fig. 8C) cells, suggesting that eIF5B depletion has a robust effect on OSCC cell migration.

To determine if eIF5B depletion from OSCC cells would affect angiogenesis, we performed the endothelial tube formation assay using the conditioned media from the control and eIF5B-depleted OSCC cells. The conditioned media from the control OSCC cells were able to effectively form endothelial tubes in HUVEC cells. However, the various parameters of endothelial tube formation, such as branching, junction formation, segments, and nodes, were significantly affected when conditioned media from eIF5B-depleted UMSCC-29 cells were used (Fig. 4E). Similarly, HUVEC cells were unable to effectively form endothelial tubes when conditioned media from eIF5B-depleted CAL-33 (Supple. Fig. 7D) or UMSCC-1 (Supple. Fig. 8D) were used. Therefore, we monitored the levels of inflammatory and pro-angiogenic cytokines in the spent media of control and eIF5B-depleted OSCC cells using a human angiogenesis and growth factor 17-plex array (Eve Technologies, University of Calgary). We observed a significant reduction in interleukin-8 (IL-8), VEGF-C, and placental growth factor (PLGF) in the conditioned media of the eIF5B-depleted UMSCC-29 cells. We also observed a significant reduction in the levels of IL-8, endothelin, PLGF, VEGFA, and VEGFC in the conditioned media of CAL-33 (Supple. Fig. 7E) and UMSCC-1 (Supple. Fig. 8E) (Supple. Table 2). However, we did not observe any significant changes in the levels of these inflammatory and pro-angiogenic cytokines in the conditioned media of eIF5B-depleted BJ-5ta immortalized fibroblast cells (Supple. Fig. 9). We further performed western blot analysis of the control and eIF5B-depleted OSCC cells to examine the levels of key drivers of invasion, migration, and proliferation. As such, we examined the effect of eIF5B depletion on EGFR, ERK, NF-κB, HIF-1α, and VEGFA expression levels. Also, the changes in phosphorylation levels of p-EGFR, p-ERK, and p-NF-κB (Fig. 4G, Supple. Figs. 7F and 8F) was assessed. Depletion of eIF5B resulted in decreased levels of total EGFR in UMSCC-29 cells (Fig. 4G). EGFR levels did not decrease in eIF5B-depleted CAL-33 (Supple. Fig. 7F) or in UMSCC-1 (Supple. Fig. 8F). However, the EGFR levels were significantly reduced in all three OSCC cell lines when eIF5B depletion was combined with TRAIL treatment. Interestingly, phosphorylation of EGRF was significantly reduced in all three OSCC cell lines under eIF5B depletion alone or combination of eIF5B depletion and TRAIL treatment (Fig. 4G, Supple. Fig. 7F, and Supple. Fig. 8F). ERK1/2 levels were significantly reduced in all three OSCC cell lines under eIF5B depletion (Fig. 4G, Supple. Fig. 7F, and Supple. Fig. 8F). However, except for CAL-33 cells, eIF5B-depleted OSCC cells did not show a significant reduction in ERK1/2 levels (Fig. 4G, Supple. Fig. 7F, and Supple. Fig. 8F). Phosphorylation of ERK was significantly decreased only in UMSCC-29 and CAL-33 cells (Fig. 4G, Supple. Fig. 7F). Phosphorylation of ERK was significantly reduced in all three OSCC cell lines when eIF5B depletion was combined with the TRAIL treatment (Fig. 4G, Supple. Fig. 7F, and Supple. Fig. 8F). The levels of total NF-kB (p65) or its phosphorylation does not seem to be greatly affected by eIF5B depletion in OSCC cells (Fig. 4G, Supple. Fig. 7F, and Supple. Fig. 8F). The levels of HIF-1α were significantly reduced in both eIF5B depletion and eIF5B depletion + TRAIL treatment in UMSCC-29 and CAL-33 cells (Fig. 4G, Supple. Fig. 7F). We could not detect HIF-1α in UMSCC-1 cells. Also, the levels of VEGFA were significantly reduced in all three OSCC cell lines under both eIF5B depletion and eIF5B depletion + TRAIL treatment (Fig. 4G, Supple. Fig. 7F, and Supple. Fig. 8F). These data suggest that eIF5B regulates proliferation, invasion and migration, and angiogenesis in OSCC cells via regulating the key drivers involved in OSCC pathophysiology.

### Translation of mRNAs encoding proteins critical for OSCC pathophysiology is reduced upon eIF5B depletion

Since eIF5B depletion altered key oncogenic processes in OSCC cells, we next investigated whether these phenotypic effects stem from impaired translation of mRNAs encoding proteins that drive tumor progression. We first confirmed shRNA-mediated depletion of eIF5B in OSCC cells (Fig. 5A, and Supple. Fig. 10A). We then performed polysome profiling experiments in eIF5B-depleted OSCC cells (Fig. 5B and Supple. Fig. 10B) and probed the polysome fractions using RT-qPCR for candidate mRNAs, including XIAP, cIAP1, Bcl-xL, VEGFA, HIF-1α, and Actin. We further determined the % mRNA distribution across the polysome fractions in control vs. eIF5B-depleted OSCC cells. The shift of the mRNAs from high to low sucrose density polysome fractions indicates the reduction in translation of these mRNAs under eIF5B depletion conditions. We observed that eIF5B depletion resulted in a robust decrease in the translation of mRNAs encoding XIAP, cIAP1, Bcl-xL, and VEGFA in UMSCC-29 cells (Fig. 5C, 5D, 5E, 5F, and 5G). However, the translation of Actin mRNA was not affected by eIF5B depletion in UMSCC-29 cells (Fig. 5H). Likewise, the translation of mRNAs encoding XIAP, cIAP1, and Bcl-xL in eIF5B-depleted CAL-33 cells (Supple Fig. 10C, 10D, and 10E) was robustly decreased. We did not observe a robust decrease in the translation of HIF1α mRNA in eIF5B-depleted either UMSCC-29 or CAL-33 cells (Fig. 5G and Supple. Fig. 10F). eIF5B depletion had a modest effect on the translation of Actin mRNA in eIF5B-depleted CAL-33 cells (Supple. Fig. 10G). Notably, the steady-state levels of these mRNAs did not decrease in eIF5B-depleted UMSCC-29 (Fig. 5I) or CAL-33 (Supple. Fig. 10). In fact, the steady-state levels of some of these mRNAs (for example, XIPA and HIF-1α) were significantly increased in eIF5B-depleted cells (Fig. 5I). Collectively, these findings suggest that eIF5B plays a critical role in OSCC pathophysiology by regulating critical factors at the mRNA translation level.

**Figure 5:**
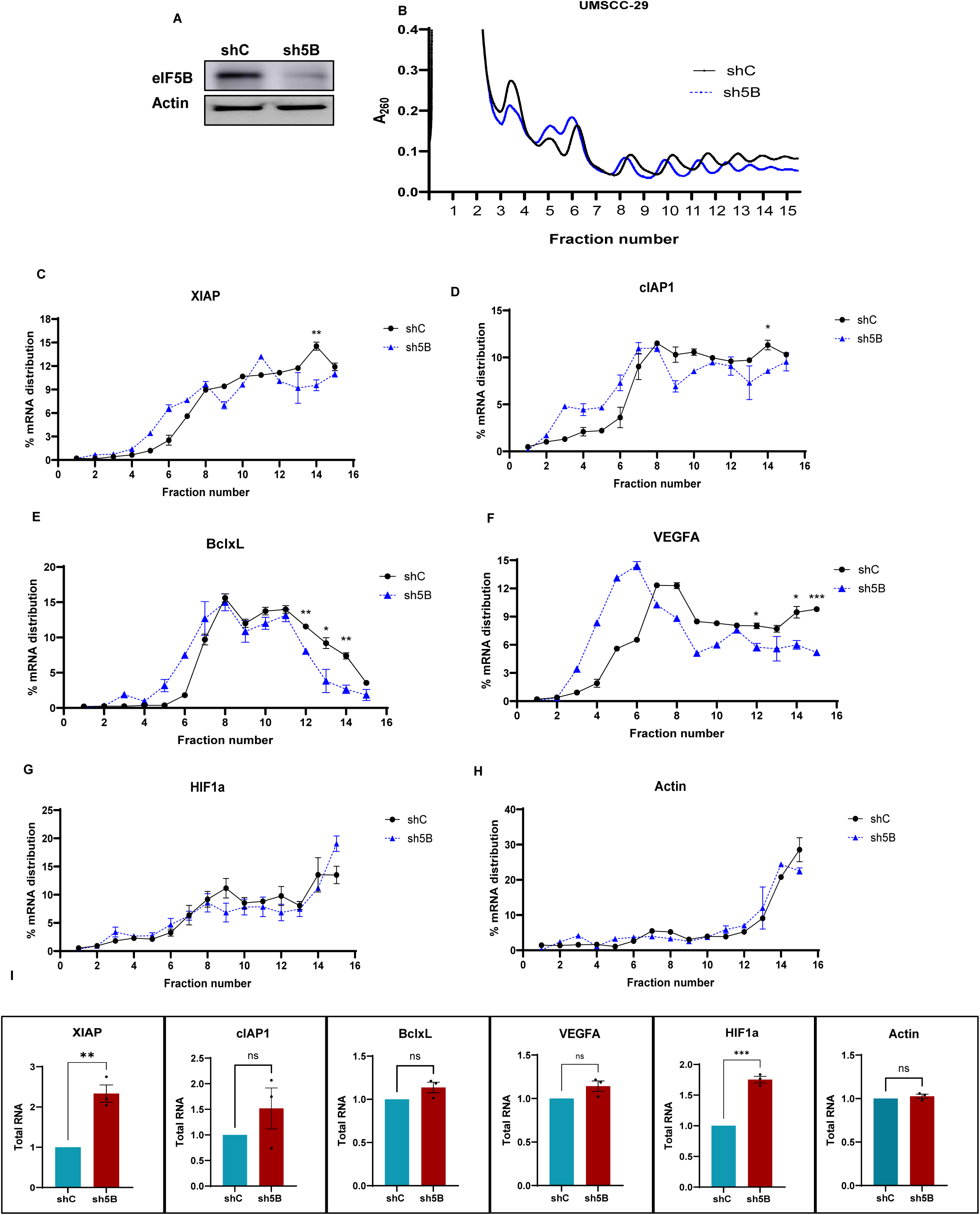
eIF5B depletion decreases the translation of distinct mRNAs encoding anti-apoptotic proteins, as well as HIF1*α,* and VEGFA. Polysome profiling analysis of *XIAP*, *Bcl-xL*, *cIAP1*, *HIF1α*, and *VEGFA* mRNAs. **(A)** Confirmation of shRNA-mediated depletion of eIF5B in UMSCC-29 cells. **(B)** Representative polysome profiles for control and eIF5B-depleted UMSCC-29 cells (black and blue curves, respectively). Total RNA was extracted from the sucrose fractions and subjected to RT-qPCR analysis. The mRNAs encoding XIAP, cIAP1, Bcl-xL, VEGFA, HIF1α, and Actin were quantified and percentage mRNA distribution across fractions was analyzed and plotted **(C, D, E, F, G, H, respectively).** Total RNA was isolated from control (shC) or eIF5B-depleted (sh5B) cells and subjected to RT-qPCR to quantify steady-state levels of the indicated mRNAs. Data are presented as mean ± SEM for three independent biological replicates. Statistical significance is indicated as *, p < 0.05; **, p < 0.01; ***, p < 0.001.

### Depletion of eIF5B impairs tumor growth and reduces the expression of critical proteins associated with OSCC pathophysiology

UMSCC-29 human oral squamous cell carcinoma (OSCC) cells were injected subcutaneously into the flank of nude mice to establish xenograft tumors (Fig. 6A). The depletion of eIF5B in implanted UMSCC-29 cells using a short hairpin RNA (sh5B) resulted in a significant reduction in tumor growth, as evidenced by significantly decreased tumor volume measurements and a visible decrease in tumor size compared with control shRNA (shC) tumors (Fig. 6B). Consistent with impaired tumor growth, western blot analysis of UMSCC-29 cells (before implantation in mice) (Fig. 6C) with eIF5B knockdown showed a significant reduction in multiple anti-apoptotic proteins such as XIAP, Bcl-xL, cIAP1, and cFLIPs. Finally, we performed immunohistochemical analysis of FFPE xenografted sections. We observed reduced DAB staining intensity in sh5B tumors compared with shC tumors for several markers, including XIAP, Bcl-xL, eIF5B, cIAP1, VEGFA, and HIF1α (Fig. 6D). In the negative control (no antibody treatment), we did not observe DAB staining (Fig. 6D, bottom panel). Moreover, we did observe DAB staining with the eIF2A antibody in both shC and sh5B xenografted tumors. As eIF2A is not the downstream target of eIF5B, it is expected that eIF2A staining does not decrease in sh5B xenografted tumors. This data supports a role for eIF5B in maintaining pro-survival and pro-angiogenic signaling in OSCC in vivo.

**Figure 6:**
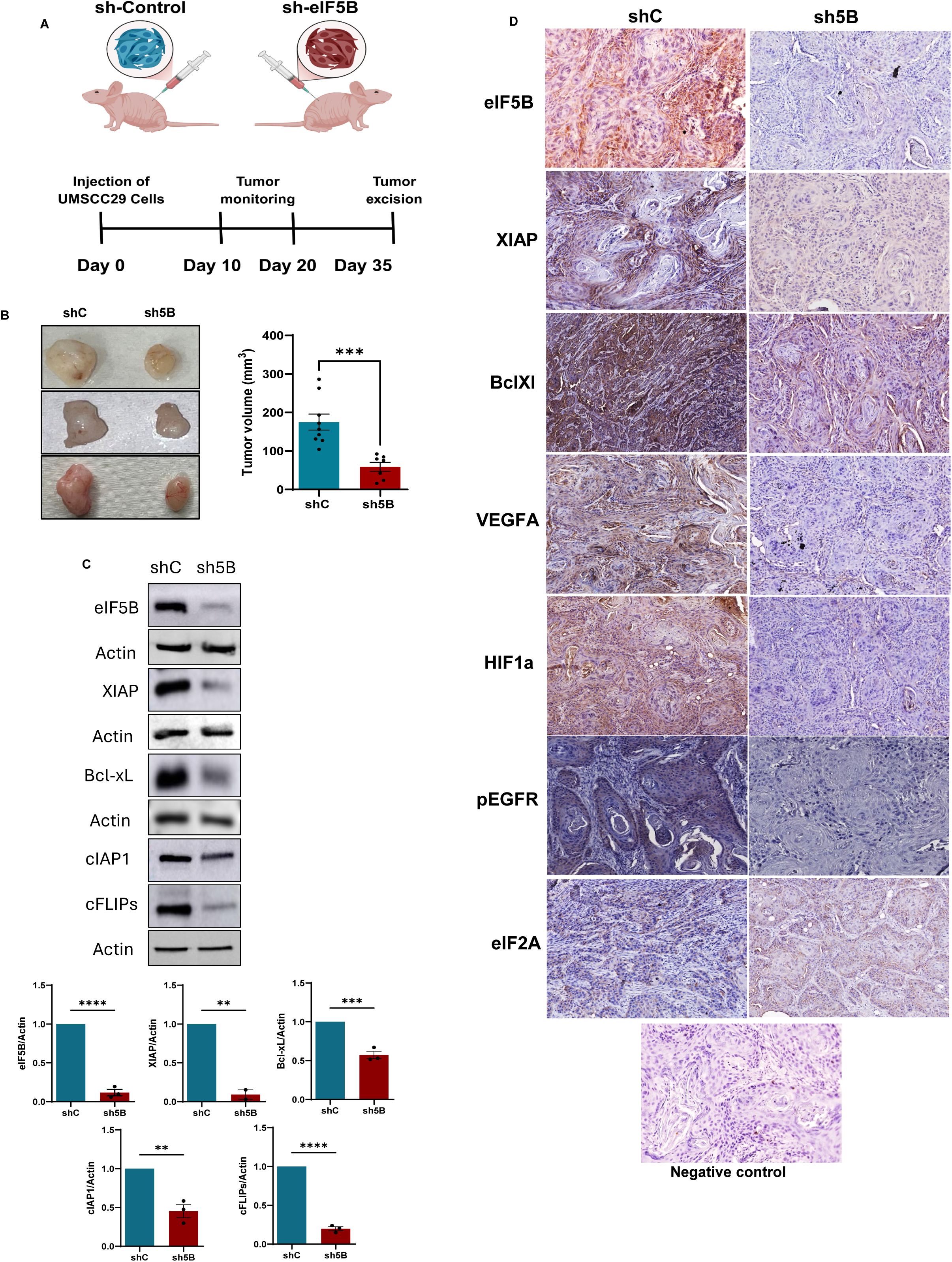
eIF5B depletion impairs tumor growth and reduces the expression of critical proteins associated with OSCC pathophysiology. **(A)** Schematic diagram of subcutaneous injection of UMSCC-29 cells (sh-Control or sh-eIF5B) in a flank model of nude mice and the tumor monitoring timeline. **(B)** Representative excised flank xenograft tumors from control and eIF5B-depleted groups at endpoint, demonstrating visible differences in tumor size. The size of tumors originated from shC (n=9)- and sh-eIF5B (n=7)-transduced UMSCC-29 cells was measured using vernier calipers. One animal from the shC group and three animals from the sh-eIF5B group did not form measurable tumors. The volume of tumors was calculated using Volume = (Length X Width^2^)/2. Data represent mean ± SEM; p < 0.001 by unpaired t-test. **(C)** Western blot analysis of shC and sh-eIF5B cells that were injected into the flank region of nude mice. Representative immunoblots for eIF5B, XIAP, Bcl-xL, cIAP1, and cFLIPs **(C, top panel)**. Data are presented as mean ± SEM from three independent biological replicates using t-test, with statistical significance indicated as follows: *, p < 0.05; **, p < 0.01; ***, p < 0.001 **(C, bottom panel)**. **(D)** The excised tumors were subjected to an immunohistochemistry assay to analyze the expression of eIF5B, XIAP, Bcl-xL, VEGFA, HIF1α, pEGFR, and eIF2A. Hematoxylin counterstain highlights nuclear architecture (blue). Representative sections are shown at [10x] magnification. Representative Hematoxylin staining of a negative control (no antibody treatment) section shows the absence of DAB-only effect **(bottom panel)**.

## Discussion

In the current study, we reveal that eIF5B is a prognostic biomarker and demonstrate that its depletion diminishes multiple cancer hallmarks, suggesting that eIF5B may represent a promising avenue for future therapeutic investigation. Analysis of genome-wide expression data from the GTEx dataset shows that eIF5B mRNA expression is significantly higher in OSCC than in matched normal oral mucosa and across all other surveyed body subsites, suggesting that eIF5B should be evaluated as a therapeutic target for OSCC treatment. We performed bioinformatic analysis of publicly available RNA-Seq and proteomics data, which showed that *EIF5B* mRNA and eIF5B protein are expressed at significantly higher levels in OSCC tumors. The single cell RNA-Seq data analysis revealed that the levels of *EIF5B* mRNA were highest in tumour cells compared to other cell types within the tumor microenvironment. Both *EIF5B* mRNA and eIF5B protein are associated with poor patient outcomes, and higher levels of EIF5B transcript is correlated with the higher lymph node metastasis of OSCC. Taken together, these data suggest that eIF5B could serve as a prognostic biomarker for OSCC tumors.

Exogenous overexpression of eIF5B in OSCC cells leads to enhanced expression of distinct antiapoptotic proteins (such as XIAP, Bcl-xL, cIAP1, and cFLIPs) (Fig. 2F, and Supple. Fig.3C). Depletion of eIF5B enhanced the TRAIL-mediated apoptosis in OSCC cells. This could be largely attributed to the decreased levels of certain anti-apoptotic proteins in eIF5B-depleted OSCC cells (Fig. 1, Supple. Fig. 1, 3, 4, 5). Despite having TRAIL receptors expressed in BJ-5ta cells, eIF5B depletion did not enhance TRAIL-induced apoptosis in BJ-5ta cells (Fig. 3). Also, the levels of distinct anti-apoptotic proteins (such as XIAP, Bcl-xL, cIAP1, and cFLIPs) were not decreased in eIF5B-depleted BJ-5ta cells. The levels of XIAP were decreased only when eIF5B depletion was combined with the treatment of an inhibitor of cap-dependent translation (Fig. 3D). This is in line with our previously published data showing that the IRES-mediated translation of XIAP was decreased only when eIF5B depletion was combined with the treatment of poly-I:C, which inhibits the cap-dependent translation, in HEK293T cells ^32^. We and others have shown that eIF5B is dispensable in non-cancer cells under basal growth conditions ^19,33,34^. To understand why cancer cells rely heavily on eIF5B for their survival, we performed an immunoprecipitation experiment for eIF5B and probed for initiator tRNA using RT-qPCR. Our data demonstrated that in OSCC cells the interaction of eIF5B with initiator tRNA is several folds higher compared to non-cancer BJ-5ta cells (Fig. 3E). This could be attributed to one or more of the following scenarios: 1) OSCC cells express higher levels of eIF5B, 2) OSCC cells express higher levels of initiator tRNA, or 3) the levels of both eIF5B and initiator tRNA are higher in OSCC cells showing higher initiator tRNA pull down with eIF5B. Importantly, Ho. et al. demonstrated that the affinity of eIF5B with initiator tRNA increases under hypoxic stress conditions in GBM cells ^33^. It is also possible that the affinity of eIF5B with initiator tRNA in OSCC cells is inherently higher compared to non-cancer BJ-5ta cells. Accordingly, OSCC cells are addicted to eIF5B for their survival, and enhanced expression of eIF5B allows cancer cells to resist apoptotic cell death. As eIF5B depletion primarily affects the pathophysiology of OSCC cells and not the non-cancer cells, targeting eIF5B using small molecule compounds in future studies may provide a therapeutic window for OSCC treatment.

Like eIF5B, eIF2A (not to be confused with eIF2α) is reported to interact with the initiator tRNA and to regulate mRNA translation ^31,35^. Therefore, we checked if eIF2A plays any role in regulating the expression of distinct antiapoptotic proteins and other markers associated with proliferation, invasion, migration, and angiogenesis. The levels of these proteins did not decrease in eIF2A-depleted OSCC cells (Supple. Fig. 6). These data suggest that although both eIF2A and eIF5B interact with initiator tRNA, only eIF5B plays a critical role in the expression of distinct anti-apoptotic proteins and other makers associated with OSCC pathophysiology. Depletion of eIF5B in OSCC cells decreased the levels of EGFR and blunted the EGFR/ERK axis. However, unlike GBM cells ^19^, the pro-inflammatory NF-kB signaling does not seem to be affected by eIF5B depletion (Fig. 4, Supple. Fig. 7, 8) in OSCC cells. The role of EGFR/ERK and NF-kB axes is well established in the survival, proliferation, and progression of OSCC ^2,36,37^. Further, the depletion of eIF5B in hepatocellular carcinoma cells results in the inhibition of EGFR- ArfGAP with SH3 domain, ankyrin repeat, and PH domain 1 (ASAP1), and EGFR-MAPK signaling pathways ^38^. The decrease in proliferation, invasion, and migration in eIF5B-depleted OSCC cells can in part be attributed to the decreased levels of p-EGFR, and p-ERK (Fig. 4, Supple. Fig. 7, 8). The proliferation of eIF5B-depleted BJ-5ta cells was modestly but significantly increased (Fig. 4B). Furthermore, the levels of pro-angiogenic cytokines (such as IL-8, Endothelin-1, PLGF, and VEGF-A/C) were robustly reduced in the conditioned media of eIF5B-depleted OSCC cells. Also, the expression of VEGF-A and HIF-1α was decreased in eIF5B-depleted OSCC cells (Fig. 4, Supple. Fig. 7, 8). The VEGF-A/HIF-1α axis is implicated in tumor angiogenesis ^39,40^. We observed that endothelial tube formation was significantly reduced when we used the conditioned media of eIF5B-depleted OSCC cells (Fig. 4, Supple. Fig. 7, 8). This data suggests that by affecting the expression of pro-angiogenic cytokines and the VEGF-A/HIF-1α axis, eIF5B would regulate the angiogenesis of OSCC tumors. Collectively, our data demonstrate that by regulating critical factors associated with OSCC pathophysiology, eIF5B would affect several cancer hallmarks, such as proliferative signaling, resistance to cell death, tumour vascularization/angiogenesis, as well as invasion and metastasis ^41^.

Under pathophysiological and physiological stress conditions, eIF5B forms an alternative ternary complex (eIF5B-GTP-Met-tRNAi^Met^) ^17,18^. Thereby, eIF5B can deliver initiator tRNA into the P site of the ribosome during IRES-mediated translation initiation. ^5,6^ As mentioned above, the interaction of eIF5B with initiator tRNA is several-fold higher in OSCC compared to BJ-5ta non-cancer cells (Fig. 3E). We have previously shown that in GBM, eIF5B regulates the translation of IRES-containing mRNAs, such as XIAP, Bcl-xL, cIAP1, and cFLIPs ^19,42^. Also in this study, we observed that the depletion of eIF5B leads to decreased levels of XIAP, Bcl-xL, and cFLIPs. The levels of cIAP1 were decreased in two OSCC cell lines (out of three tested) when eIF5B was depleted (Fig. 2, Supple. Fig. 3, 4). Therefore, we hypothesized that, like in GBM cells, eIF5B would regulate the translation of IRES-containing mRNAs. To this end, we performed a polysome profiling experiment and probed for selected IRES-containing mRNA in polysome fractions (Fig. 5, Supple. Fig. 10). We demonstrated that the translation of mRNAs encoding anti-apoptotic proteins, such as XIAP, Bcl-xL, and cIAP1 was reduced in both UMCC29 and CAL-33 cells. These data suggest that by regulating the translation of mRNA encoding distinct anti-apoptotic proteins, eIF5B regulates apoptosis in OSCC cells. Notably, expression of HIF-1α and VEGF-A is regulated at the translational level via IRES elements ^43,44^. Accordingly, it is very likely that eIF5B facilitates angiogenesis in OSCC by regulating the translation of mRNAs encoding key components such as VEGF-A, and HIF-1α. Indeed, depletion of eIF5B in OSCC cells resulted in a robust decrease of translation of VEGF-A mRNA (Fig. 5). Although the translation of HIF-1α mRNA is known to be regulated via an IRES element, we did not observe a major decrease in its translation in the eIF5B-depleted OSCC cells (Fig. 5 and Supple. Fig. 10). The translation of mRNA encoding Actin was not affected by the depletion of eIF5B. Also, the steady-state levels of these mRNA were not decreased. Collectively, these data suggest that eIF5B is a regulator of translation of IRES-containing mRNAs, driving several biological processes involved in OSCC pathophysiology. Interestingly, immunohistochemistry analysis of the mouse xenografted tumors demonstrates that the levels of IRES-controlled proteins (such as XIAP, Bcl-xL, VEGFA, and HIF1α) were decreased when eIF5B-depleted UMSCC-29 cells were grafted in mice (Fig. 6). Moreover, the levels of phosphorylated EGFR were decreased in eIF5B-depleted tumors (Fig. 6). Also, the depletion of eIF5B in UMSCC-29 cells significantly reduced OSCC tumor formation in the flank xenograft model (Fig. 6), suggesting that eIF5B, via modulating the critical IRES-regulated proteins, impacts the pathophysiology of OSCC tumors. Importantly, due to the critical clinical challenges associated with current standard-of-care OSCC treatments, including high morbidity and poor patient outcomes, there is an urgent need to identify new therapeutic targets. Moreover, there is a limited number of prognostic biomarkers available for the diagnosis of OSCC. In the present study, we have demonstrated that eIF5B is a prognostic biomarker and emerging therapeutic target for OSCC treatment.

## Acknowledgments

This work was funded by a Cancer Research Society- Operating Grant (1280259). Natural Science and Engineering aspect of this research was funded by the Natural Sciences and Engineering Research Council of Canada-Discovery Grants (RGPIN-2017-05463 and RGPIN-2024-04035), and the Canada Foundation for Innovation-John R. Evans Leaders Fund (35017). Pavan Narasimha and Veda Hegde were supported by the Alberta Innovates Graduate Fellowship. Jinay Patel was partially supported by the NSERC-CREATE grant (510937-2018). Beruwalage Fernando and Veda Hegde were partially supported by the NSERC-CREATE grant (584836-2024).

## Author Contributions

Conceptualization, NT, PB, SMJ, PLN; Methodology, PLN, JP, AC, VH, BF, HS, RM, AS, SCN, BYA; Analysis, PLN, JP, AC, TWM, SC, RH, JCD, MH, ET; Investigation, PLN, JP, AC; Resources, NT, PB; Writing – Original Draft, PLN, NT, PB, AC, SMJ; Writing – Review and Editing, PLN, NT, PB, AC, SMJ; Supervision, NT, SMJ, PB; Funding Acquisition, NT, PB

## Supplementary figures

**Supplementary Figure 1:**
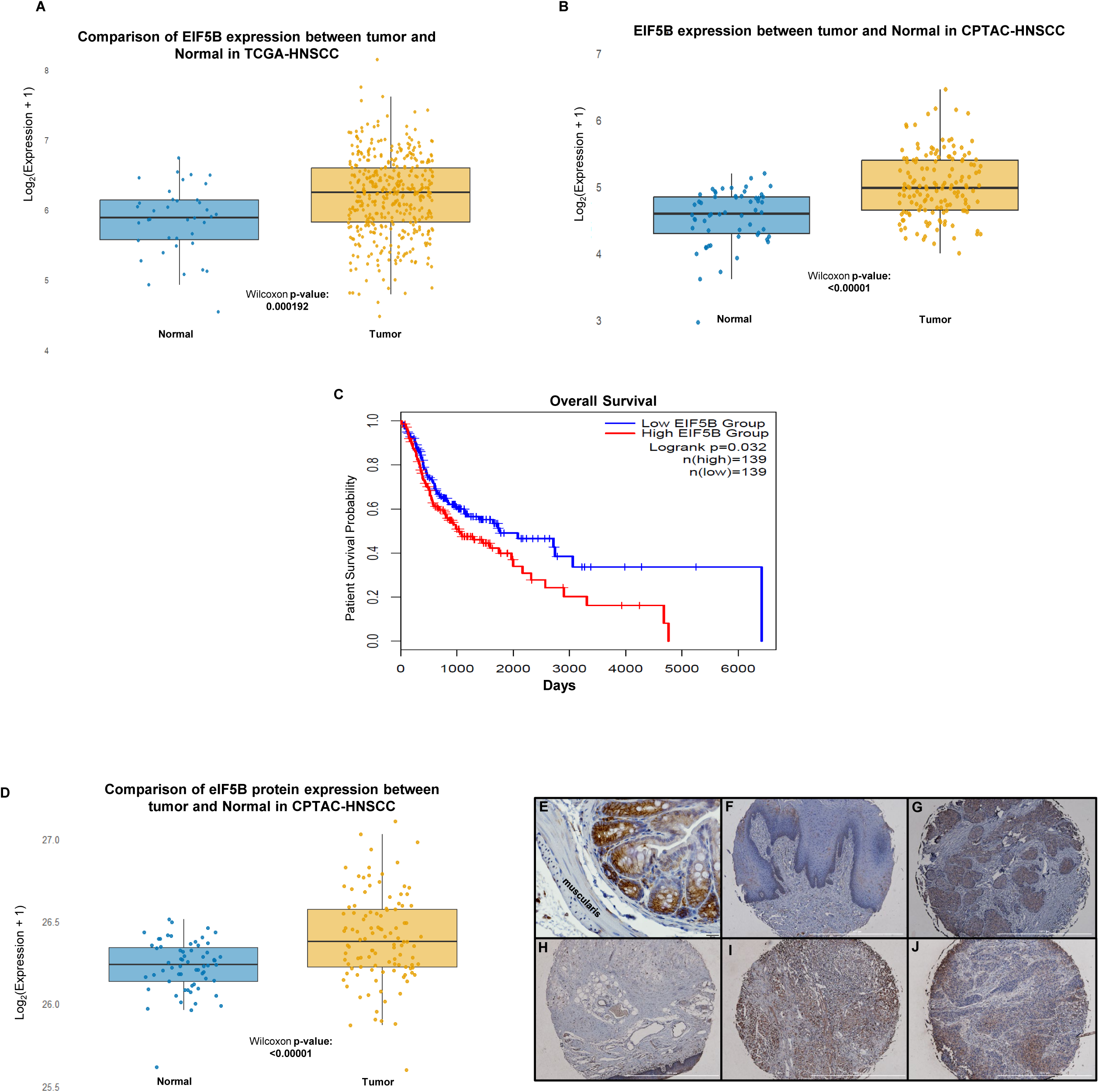
HNSCC tumors express high levels of eIF5B compared to normal samples. **(A)** *EIF5B* mRNA expression in HNSCC tumors and normal samples. Boxplot showing log2(expression + 1) of *EIF5B* mRNA in normal (blue) and tumor (yellow) samples from TCGA-HNSCC (The Cancer Genome Atlas – Head and Neck Squamous Cell Carcinoma) HPV-negative cohort (Wilcoxon test, p <0.001). **(B)** Boxplot showing log2(expression + 1) of *EIF5B* mRNA in normal (blue) and HNSCC tumor (yellow) samples from the CPTAC-HNSCC cohort (Wilcoxon test, p = 0.000192). **(C)** TCGA RNAseq data analysis shows that higher EIF5B (transcript) expressing HNSCC tumors have significantly (p = 0.032) lower patient survival times. **(D)** Box plot showing log2(expression + 1) of eIF5B protein in normal (blue) and HNSCC tumor (yellow) samples from the CPTAC-HNSCC cohort (Wilcoxon test, p = 0.000192). **(E)** We have optimized immunohistochemical-staining conditions for eIF5B. Smooth muscle (muscularis) expresses very little eIF5B. Therefore, we used the mouse colon as a negative control. This antibody can react with both mouse and human eIF5B. The expression of eIF5B in the muscularis is almost null. **(F-J)** We also tested the eIF5B antibody using commercially available TMAs (34 normal and 180 HNSCC tissues), Catalogue # HN118 and OR601c (https://www.tissuearray.com). We compared the expression of eIF5B in cancer adjacent tongue **(H)** and cancer adjacent larynx tissue **(I)** with tongue cancer **(G and H)**, and larynx cancer tissues **(J).**

**Supplementary Figure 2:**
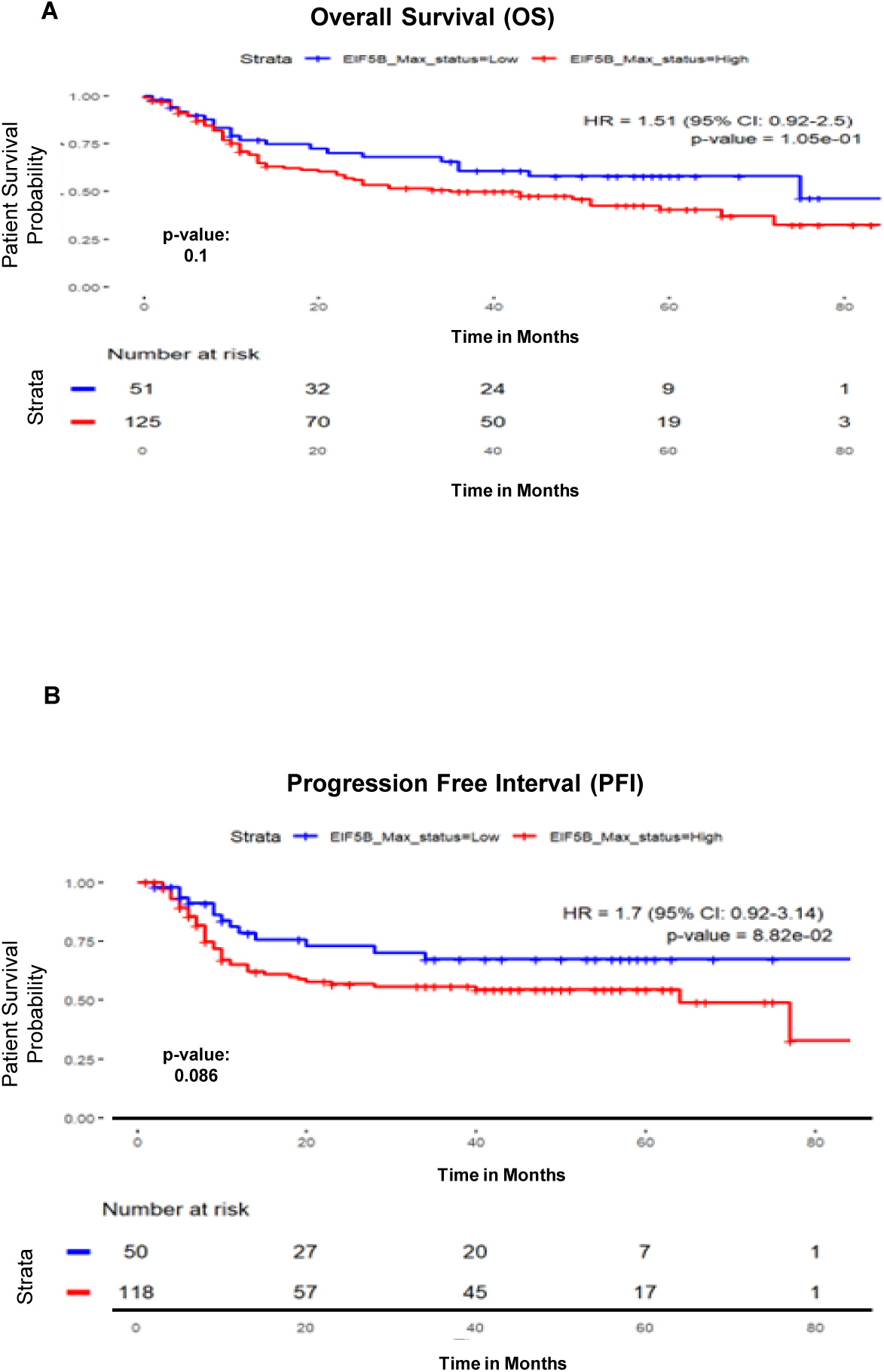
HNSCC express high levels of eIF5B, which leads to poorer survival rates. **(A)** Kaplan–Meier overall survival curves of OSCC by high (2/3) vs. low (0/1) eIF5B expression levels. (B) Kaplan–Meier progression-free survival curves of OSCC by high (2/3) vs. low (0/1) eIF5B expression levels.

**Supplementary Figure 3:**
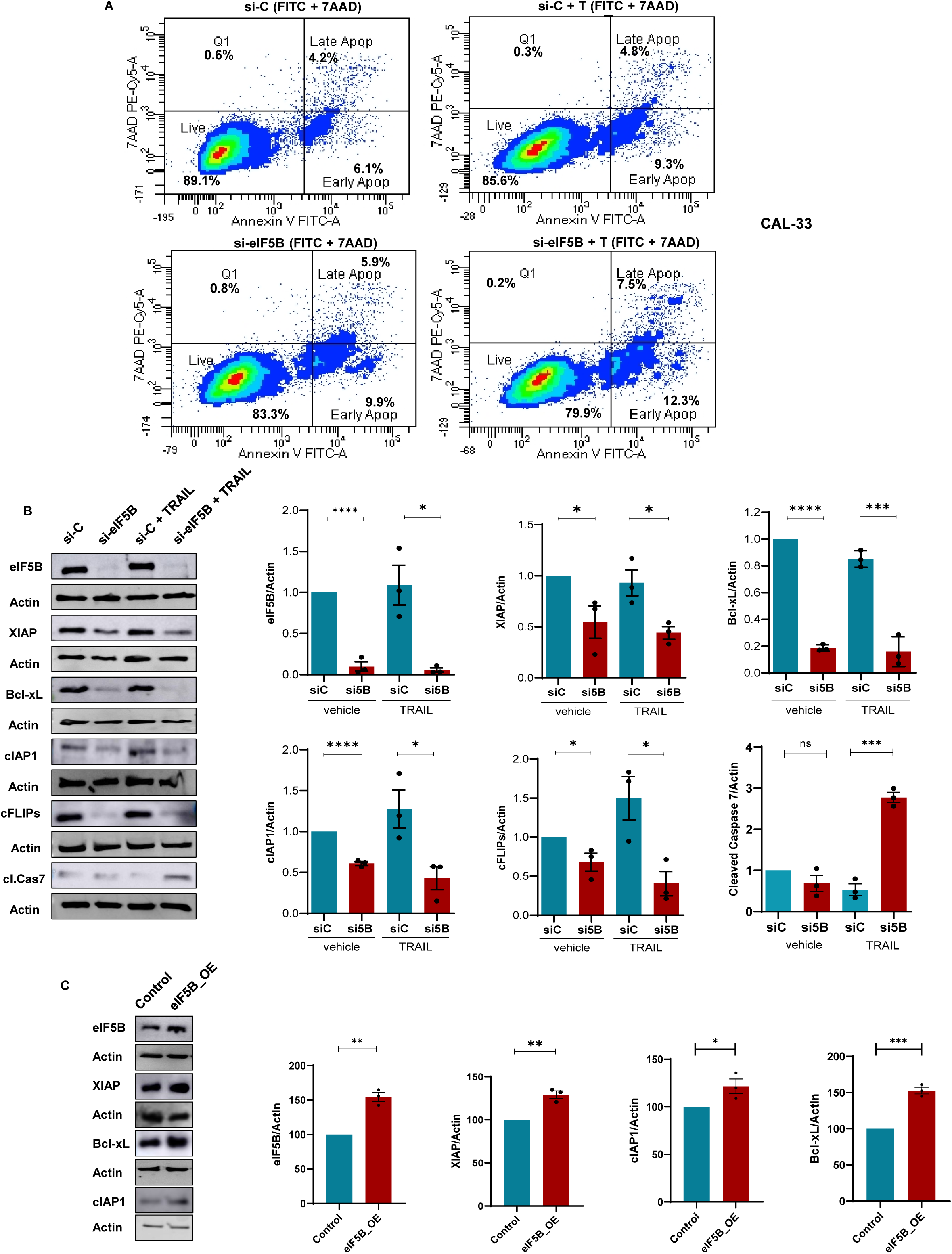
eIF5B knockdown enhances TRAIL-induced apoptosis and modulates anti-apoptotic protein expression in CAL-33. **(A)** Flow cytometry analysis of Annexin V-FITC/7-AAD staining in CAL-33 cells transfected with control (si-C) or eIF5B-targeting siRNA (si-eIF5B), treated with or without TRAIL. Quadrants denote live cells (Q3), dead cells (Q1), early apoptotic (Q4), and late apoptotic (Q2) populations. eIF5B depletion increases apoptosis (Q2: CAL-33 si5B 7.5% vs. siC 4.8% under TRAIL treatment) via annexin V and 7AAD labelling. **(B)** Western blot analysis shows eIF5B knockdown reduces anti-apoptotic proteins XIAP, Bcl-xL, cIAP1, and cFLIPs in CAL-33 and increases cleaved caspase-7 (cl.Cas7) under si-eIF5B + TRAIL treatment. Densitometry (normalized to actin) confirms significant reductions in XIAP, Bcl-xL, cIAP1, and cFLIPs under si-eIF5B as well as TRAIL-treated groups. Data are presented as mean ± SEM for three biological replicates. Statistical significance is indicated as *, p < 0.05; **, p < 0.01; ***, p < 0.001; ****, p < 0.0001

**Supplementary Figure 4:**
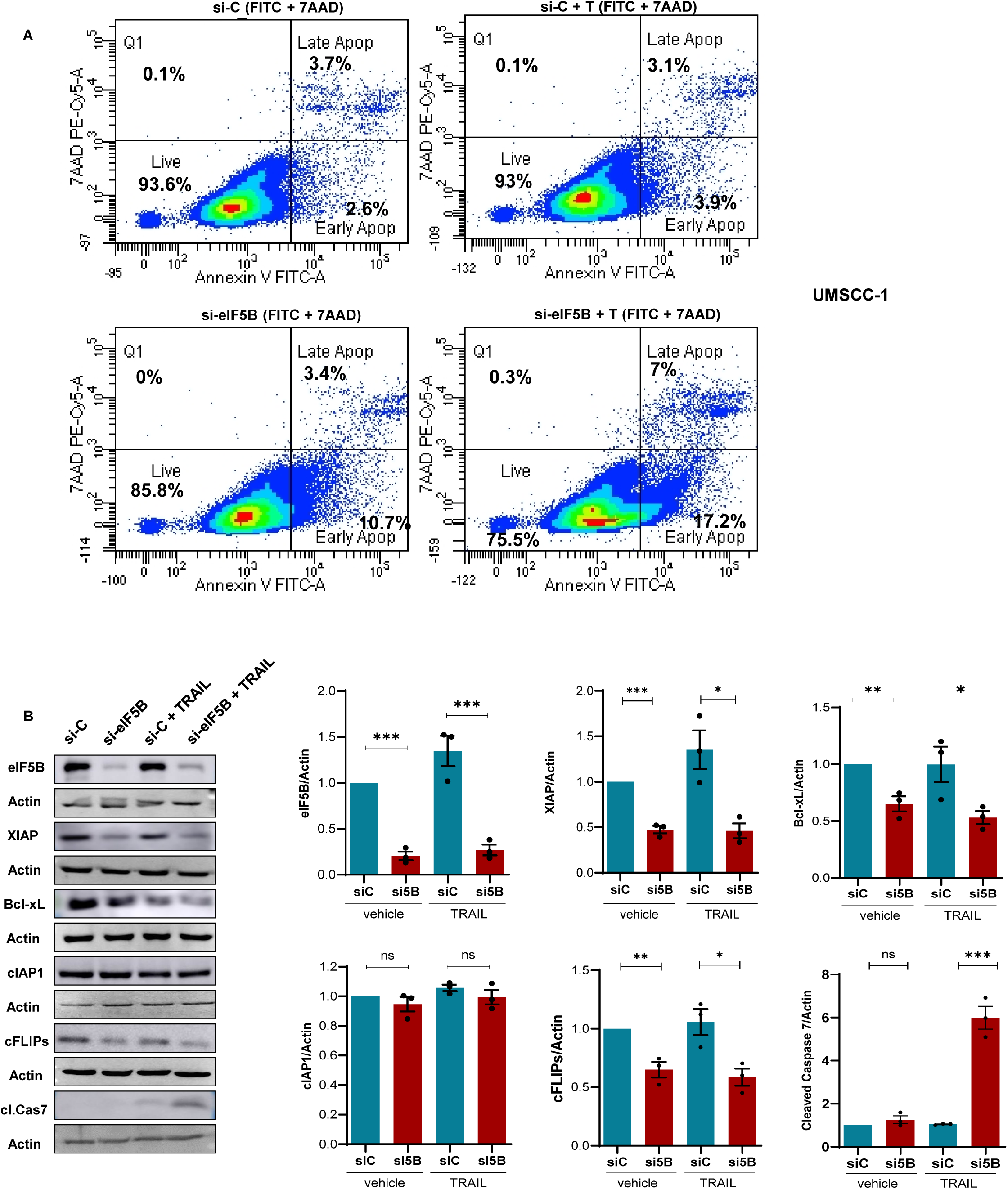
eIF5B knockdown enhances TRAIL-induced apoptosis and modulates anti-apoptotic protein expression in UMSCC-1 cells. **(A)** Flow cytometry analysis of Annexin V-FITC/7-AAD staining in UMSCC-1 cells transfected with control (si-C) or eIF5B-targeting siRNA (si-eIF5B), treated with or without TRAIL. Quadrants denote live cells (Q3), dead cells (Q1), early apoptotic (Q4), and late apoptotic (Q2) populations. eIF5B depletion increases apoptosis (Q2: UMSCC-1 si5B 7% vs. siC 3.1% with TRAIL) via annexin V and 7AAD labelling. **(B)** Western blot analysis shows eIF5B knockdown reduces anti-apoptotic proteins XIAP, Bcl-xL, and cFLIPs and increases cleaved caspase-7 (cl.Cas7) under si-eIF5B + TRAIL treatment. However, cIAP1 was not affected in UMSCC-1 cells. Densitometry (normalized to actin) confirms significant reductions in XIAP, Bcl-xL, and cFLIPs under si-eIF5B as well as TRAIL-treated groups. Data are presented as mean ± SEM for three biological replicates. Statistical significance is indicated as *, p < 0.05; **, p < 0.01; ***, p < 0.001; ****, p < 0.0001

**Supplementary Figure 5.**
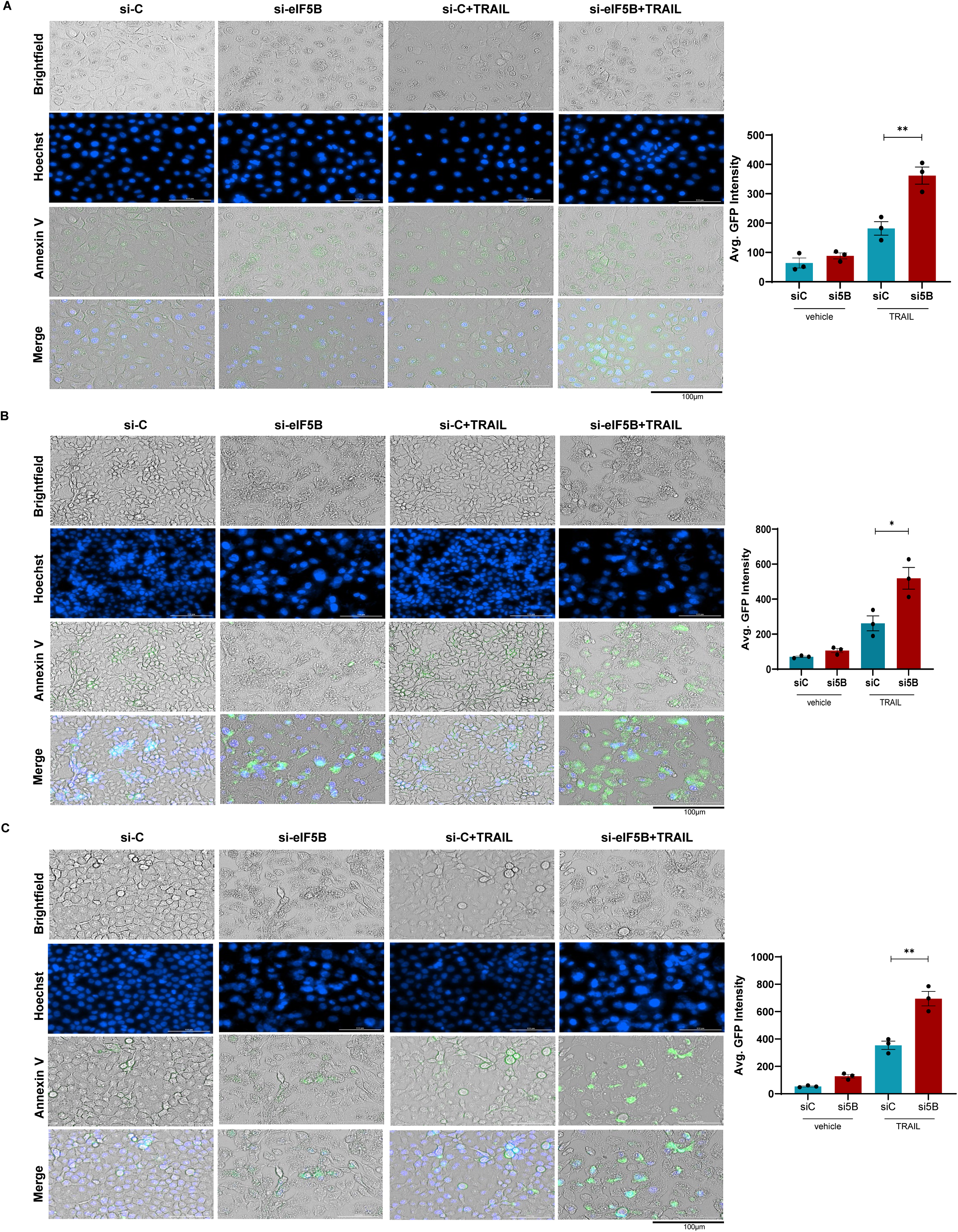
eIF5B depletion enhances TRAIL-induced apoptosis in OSCC cells. **(A-C)** Fluorescence microscopy images (Brightfield, Hoechst nuclear stain, Annexin V-FITC apoptosis marker, and merged channels) of siRNA-transfected cells treated with TRAIL (100 ng/mL). A-C: Representative images show increased Annexin V staining (green) and nuclear fragmentation (Hoechst, blue) in si-eIF5B + TRAIL groups compared to controls (si-C). Bar graphs quantify Annexin V-FITC intensity (average GFP fluorescence intensity) under si-C vs. si-eIF5B conditions with/without TRAIL. Data represent mean ± SEM (n=3 biological replicates; *p < 0.05, **p < 0.01, one-way ANOVA). Scale bars: 100 µm.

**Supplementary Figure 6:**
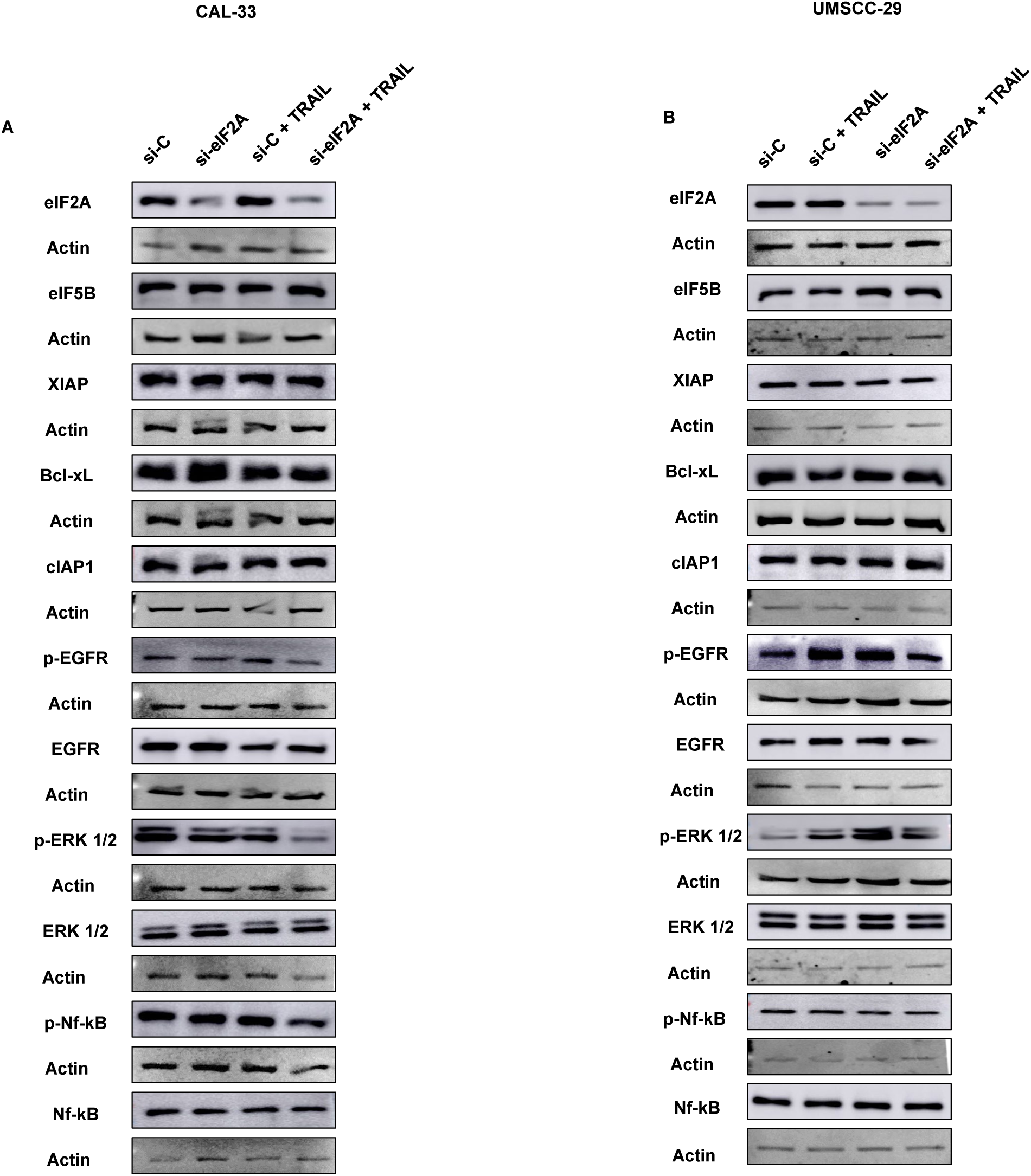
eIF5B is unique among different initiation factors. **(A & B)** Western blot analysis was performed in CAL-33 and UMSCC-29 cells transfected with control (si-C) or eIF2A-targeting siRNA (si-eIF2A), treated with or without TRAIL. The levels of XIAP, cIAP1, Bcl-xL, EGFR, ERK, and NF-kB were monitored.

**Supplementary Figure 7.**
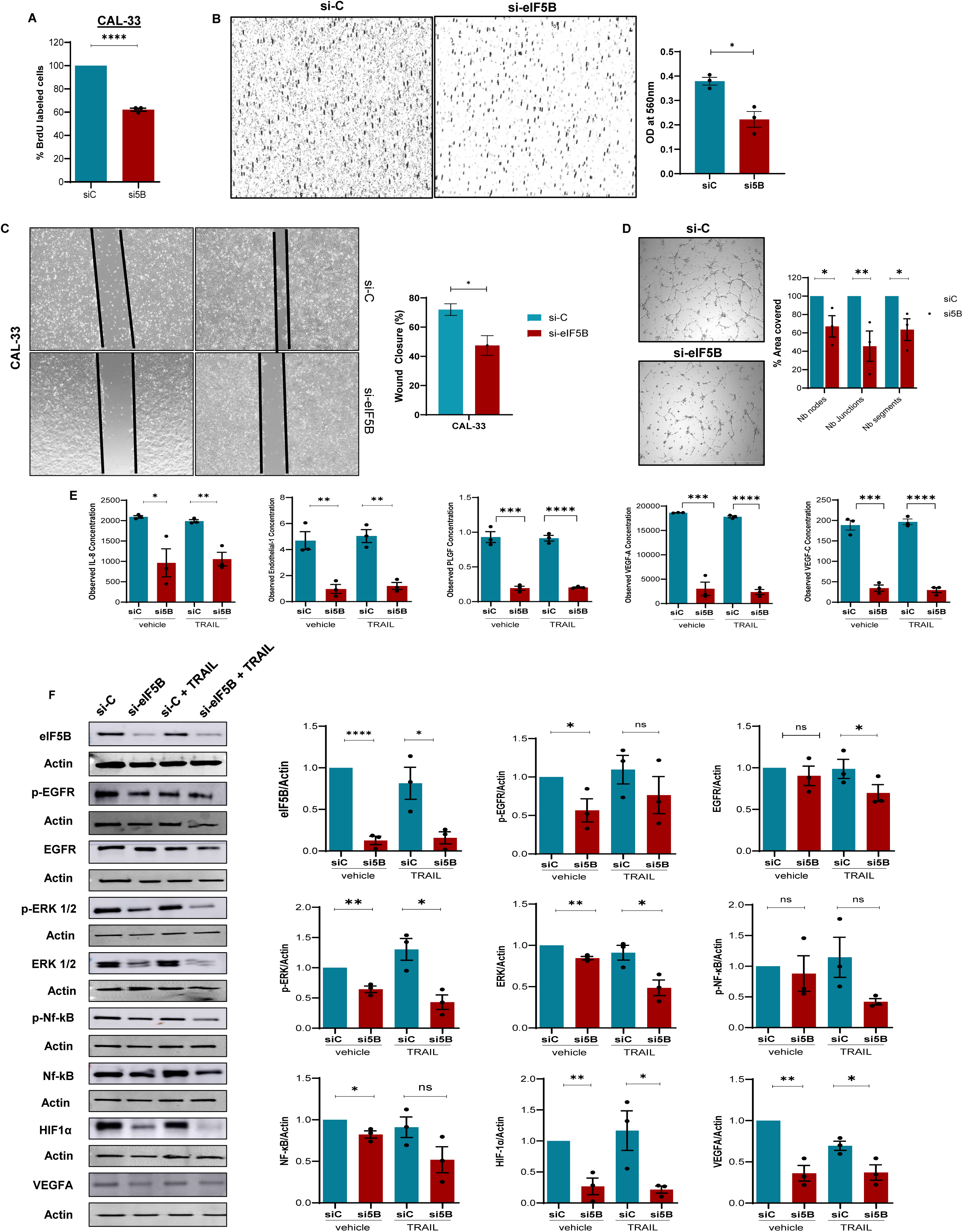
eIF5B depletion decreases the proliferation, invasion, migration and angiogenesis of CAL-33 cells. **(A)** BrdU incorporation assay was used to measure cell proliferation. Quantification shows the percentage of BrdU-labeled cells in control (siC) and si5B-treated groups. Data are presented as mean ± SEM; ***p < 0.001. **(B)** A collagen-based invasion assay was used to evaluate cell invasiveness in control (siC) and si5B-treated groups. Cells were seeded onto collagen matrices and allowed to invade for a defined period (48-72 hours). The number of invading cells was quantified and expressed as the mean ± SEM. Microscopy images of invading cells **(B; left panel)**, quantification **(B; right panel)**. **(C)** Phase contrast microscopy images showing wound area in siC/si5B-treated CAL-33 cells at the time of scratch introduction (T = 0 h) & at T = 30 h wound closure **(C; left panel)**. Quantification of the percentage of wound area covered after 30 hours, normalized to wound area at the time of scratch introduction **(C; right panel)**. **(D)** Representative images of the endothelial HUVEC cells tube formation assay treated with conditioned media from siC- or si5B-transfected CAL-33. ImageJ angiogenesis macro software quantified Junctions, segments, and nodes. CAL-33 cells were transfected with either control siRNA (siC) or eIF5B-specific siRNA (si5B) and incubated for 96 hours. **(E)** The conditioned media for the control and eIF5B-depleted CAL-33 cells were analyzed for angiogenic biomarkers **(lower panel)** using EVE Technologies. **(G)** Representative images of immunoblots quantified probing for eIF5B, p-EGFR, EGFR, p-ERK, ERK, p-NF-kB, NF-kB, HIF1α, VEGFA, and β-actin (internal control). Data are presented as mean ± SEM from three independent biological replicates, with statistical significance indicated as follows: *, p < 0.05; **, p < 0.01; ***, p < 0.001, and ****, p< 0.0001. Scale bars: 100 µm.

**Supplementary Figure 8.**
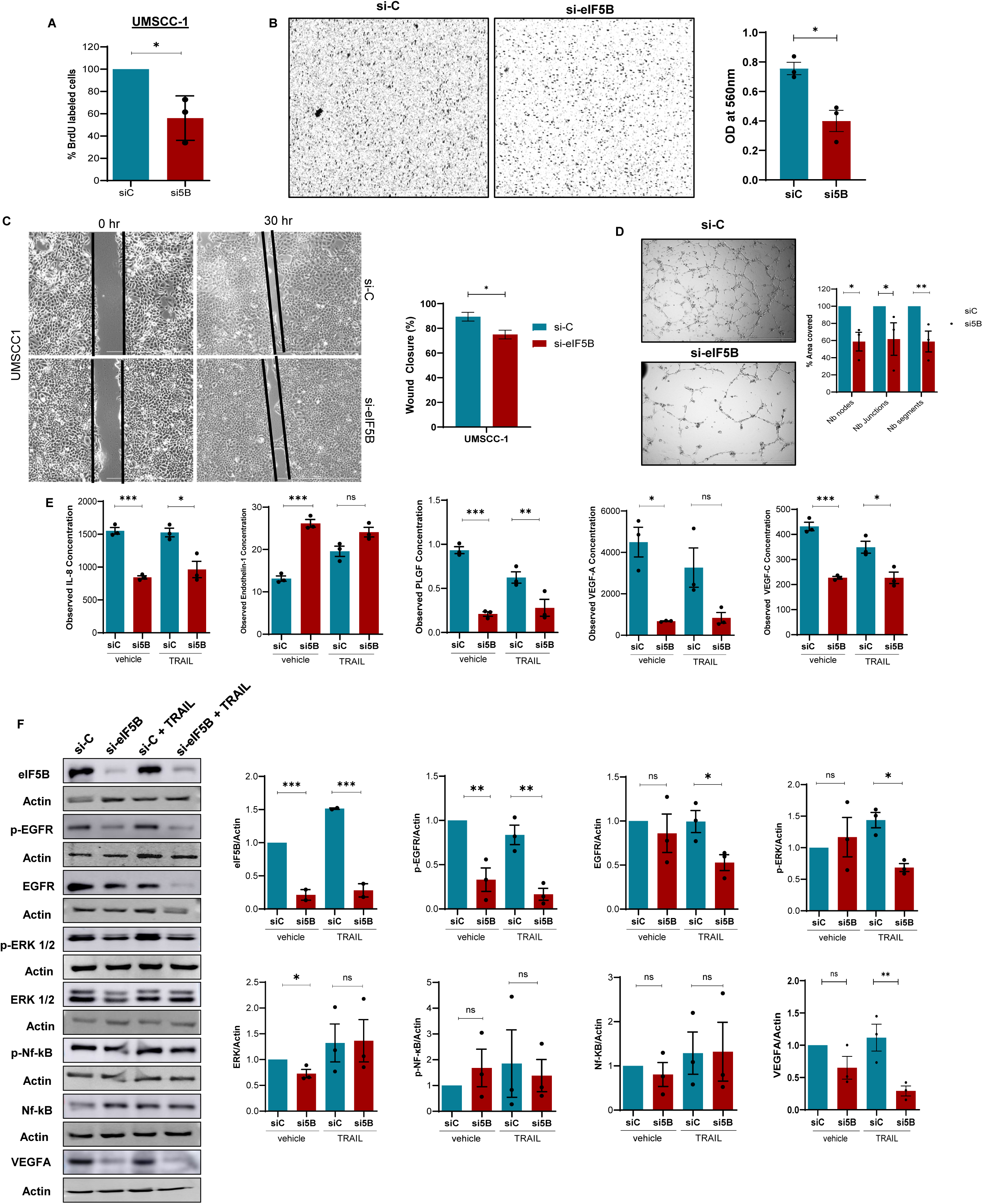
eIF5B depletion decreases the proliferation, invasion, migration and angiogenesis of UMSCC-1 cells. **(A)** BrdU incorporation assay was used to measure cell proliferation. Quantification shows the percentage of BrdU-labeled cells in control (siC) and si5B-treated groups. Data are presented as mean ± SEM; ***p < 0.001. **(B)** A collagen-based invasion assay was used to evaluate cell invasiveness in control (siC) and si5B-treated groups. Cells were seeded onto collagen matrices and allowed to invade for a defined period (48-72 hours). The number of invading cells was quantified and expressed as the mean ± SEM. Microscopy images of invading cells **(B; left panel)**, quantification **(B; right panel)**. **(C)** Phase contrast microscopy images showing wound area in siC/si5B-treated CAL-33 cells at the time of scratch introduction (T = 0 h) & at T = 30 h wound closure **(C; left panel)**. Quantification of the percentage of wound area covered after 30 hours, normalized to wound area at the time of scratch introduction **(C; right panel)**. **(D)** Representative images of the endothelial HUVEC cells tube formation assay treated with conditioned media from siC- or si5B-transfected UMSCC-1. ImageJ angiogenesis macro software quantified Junctions, segments, and nodes. UMSCC-1 cells were transfected with either control siRNA (siC) or eIF5B-specific siRNA (si5B) and incubated for 96 hours. **(E)** The conditioned media for the control and eIF5B-depleted UMSCC-1 cells were analyzed for angiogenic biomarkers **(lower panel)** using EVE Technologies. **(G)** Representative images of immunoblots quantified probing for eIF5B, p-EGFR, EGFR, p-ERK, ERK, p-NF-kB, NF-kB, HIF1α, VEGFA, and β-actin (internal control). Data are presented as mean ± SEM from three independent biological replicates, with statistical significance indicated as follows: *, p < 0.05; **, p < 0.01; ***, p < 0.001, and ****, p< 0.0001. Scale bars: 100 µm.

**Supplementary Figure 9:**
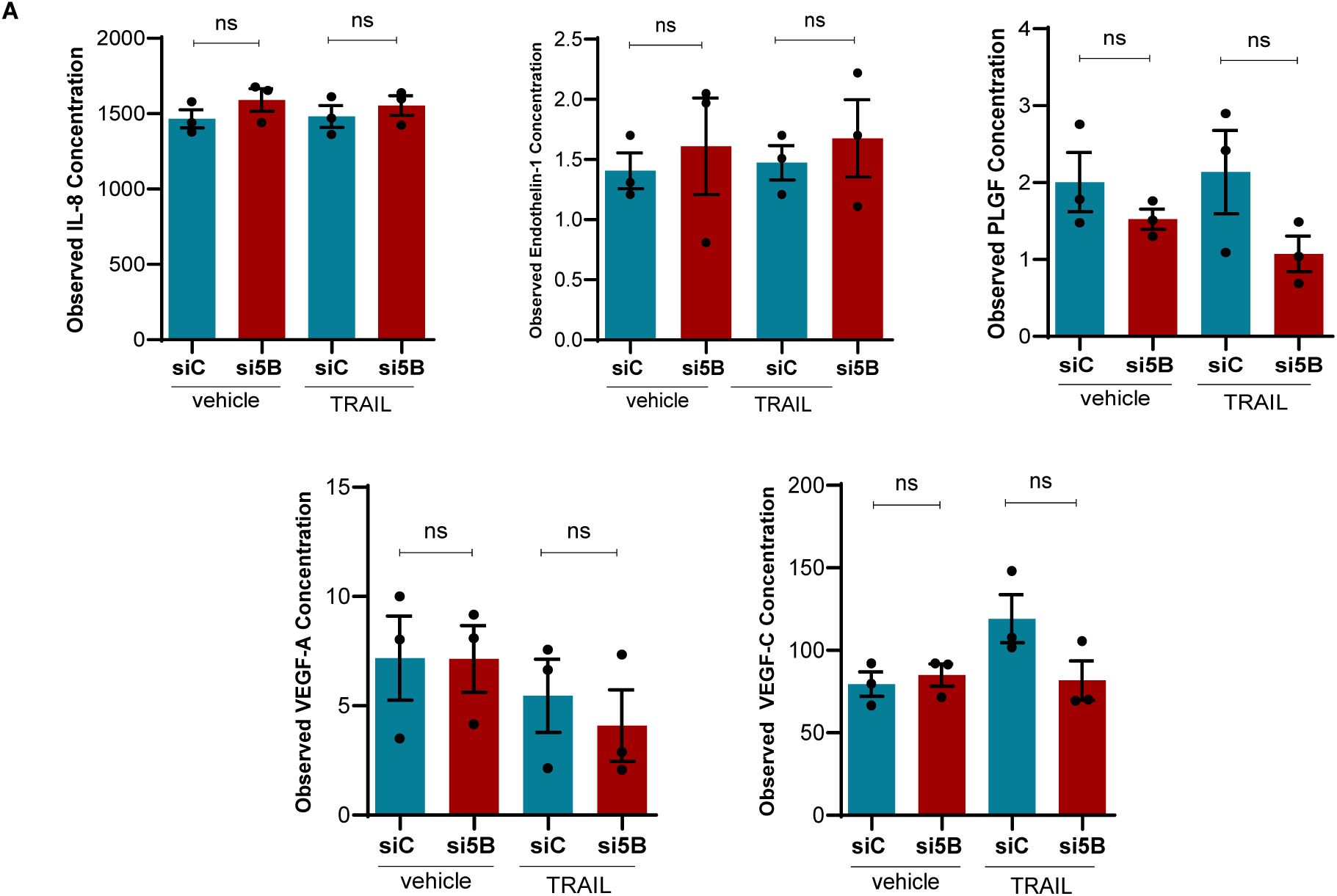
The depletion of eIF5B in BJ-5Ta (non-cancer fibroblast) cells did not significantly decrease the levels of angiogenic biomarkers. BJ-5ta cells were transfected with either control siRNA (siC) or eIF5B-specific siRNA (si5B) and incubated for 96 hours. TRAIL (100 ng/mL) treatment was performed for 4 hours, then the media was collected. The conditioned media for the control and eIF5B-depleted BT5Ta cells were analyzed for angiogenic biomarkers using EVE Technologies. The level of distinct angiogenic biomarkers was decreased in the spent media of eIF5B-depleted BJ-5ta cells. Data are expressed as mean ± SEM for three biological replicates. Statistical significance is indicated as *, p < 0.05; **, p < 0.01; ***, p < 0.001; ****, p < 0.0001.

**Supplementary Figure 10:**
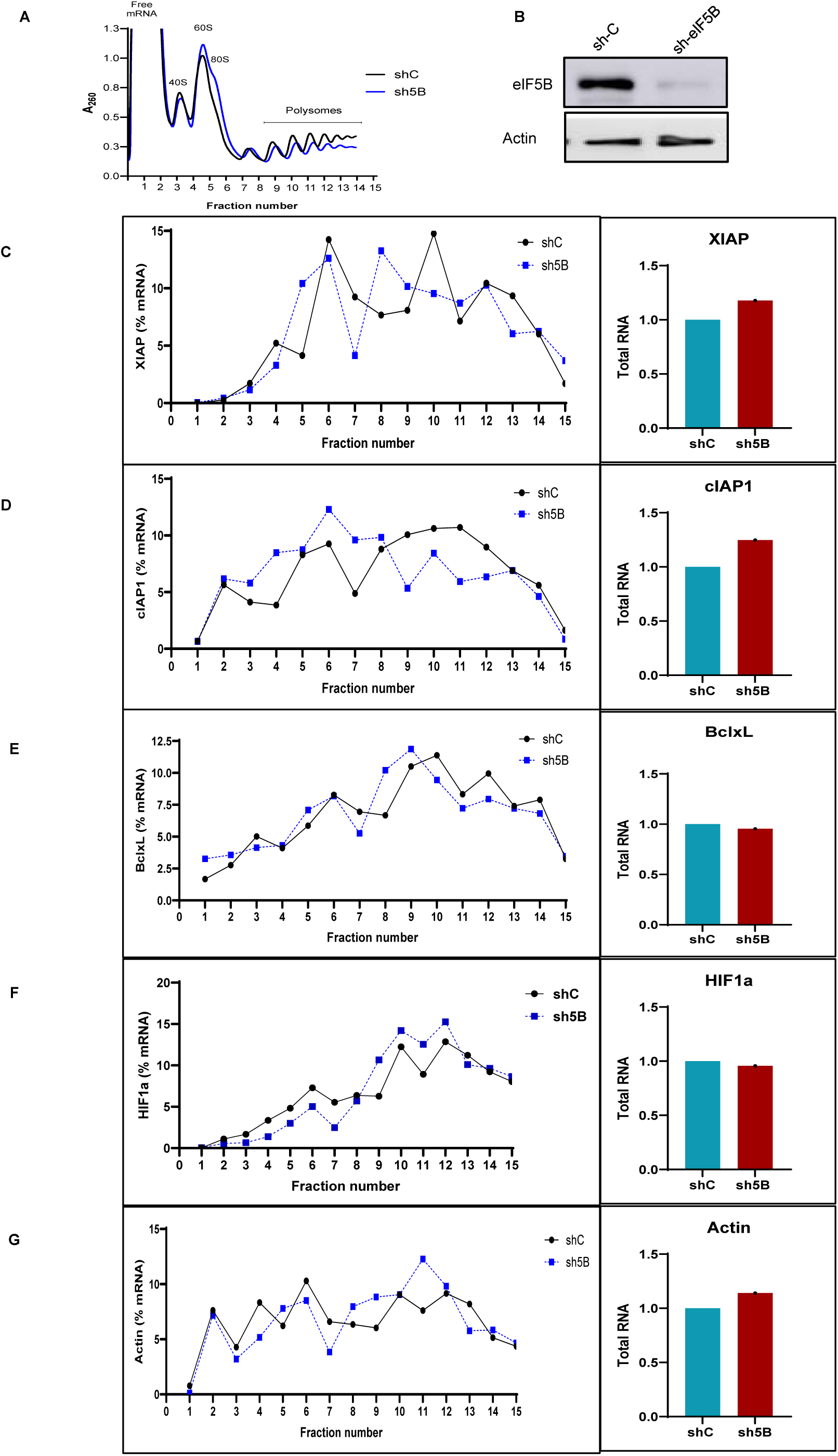
eIF5B depletion decreases the translation of distinct mRNAs encoding anti-apoptotic proteins and HIF1*α*. Polysome profiling analysis of XIAP, Bcl-xL, cIAP1, and HIF1α, mRNAs. **(A)** Confirmation of shRNA-mediated depletion of eIF5B in CAL-33 cells. **(B)** Representative polysome profiles for control and eIF5B-depleted CAL-33 cells (black and blue curves, respectively). Total RNA was extracted from the sucrose fractions and subjected to RT-qPCR analysis. The levels of mRNAs encoding XIAP, cIAP1, Bcl-xL, HIF1α, and Actin were quantified and percentage mRNA distribution across fractions was analyzed and plotted **(C, D, E, F; left panels, respectively).** Total RNA was isolated from control (shC) or eIF5B-depleted (sh5B) cells and subjected to RT-qPCR to quantify steady-state levels of the indicated mRNAs, normalized to β-actin mRNA **(C, D, E, F; right panels, respectively)**. Data are presented as mean ± SEM for three independent biological replicates. Statistical significance is indicated as *, p < 0.05; **, p < 0.01; ***, p < 0.001.

---

**Supplementary Table 1.**
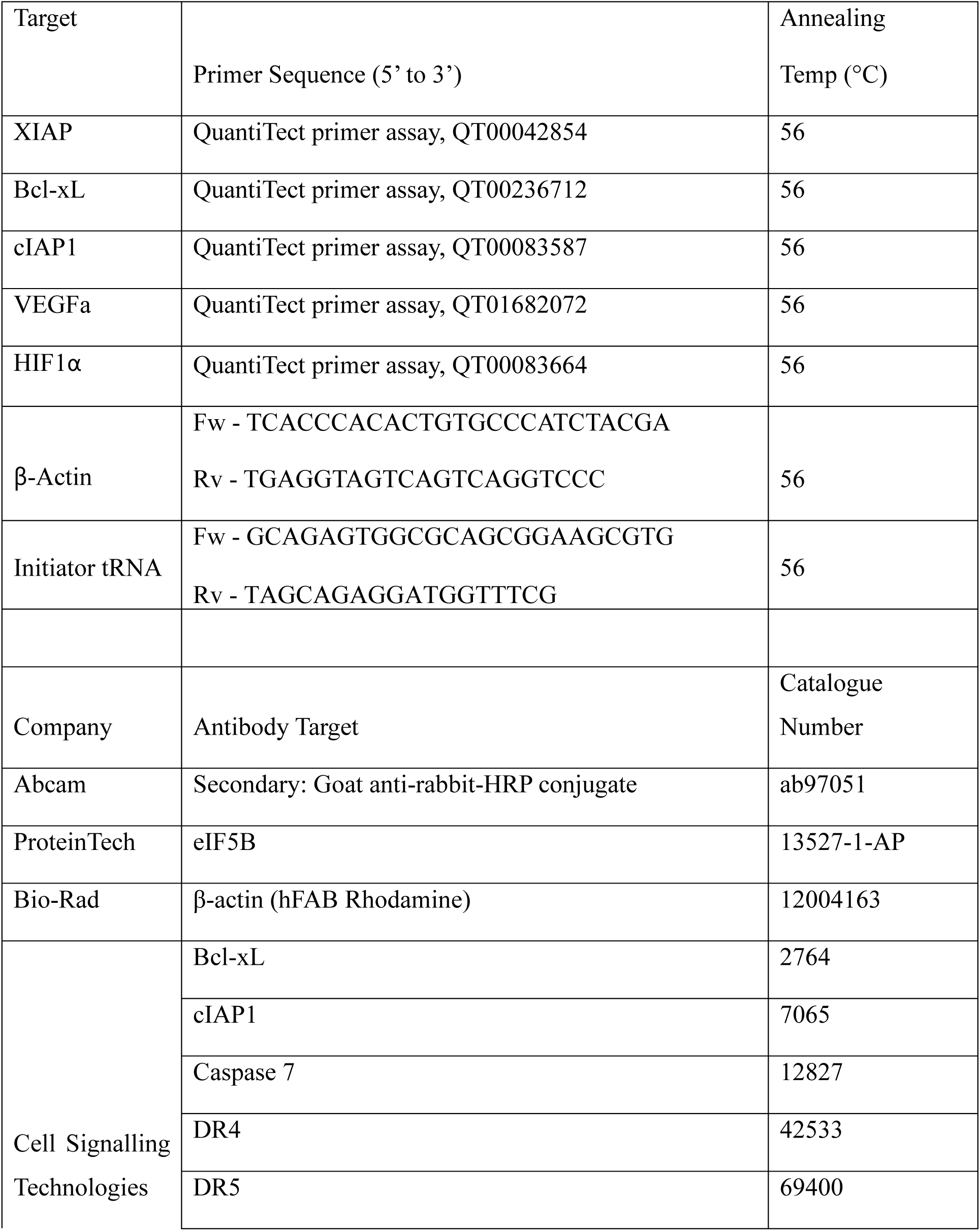
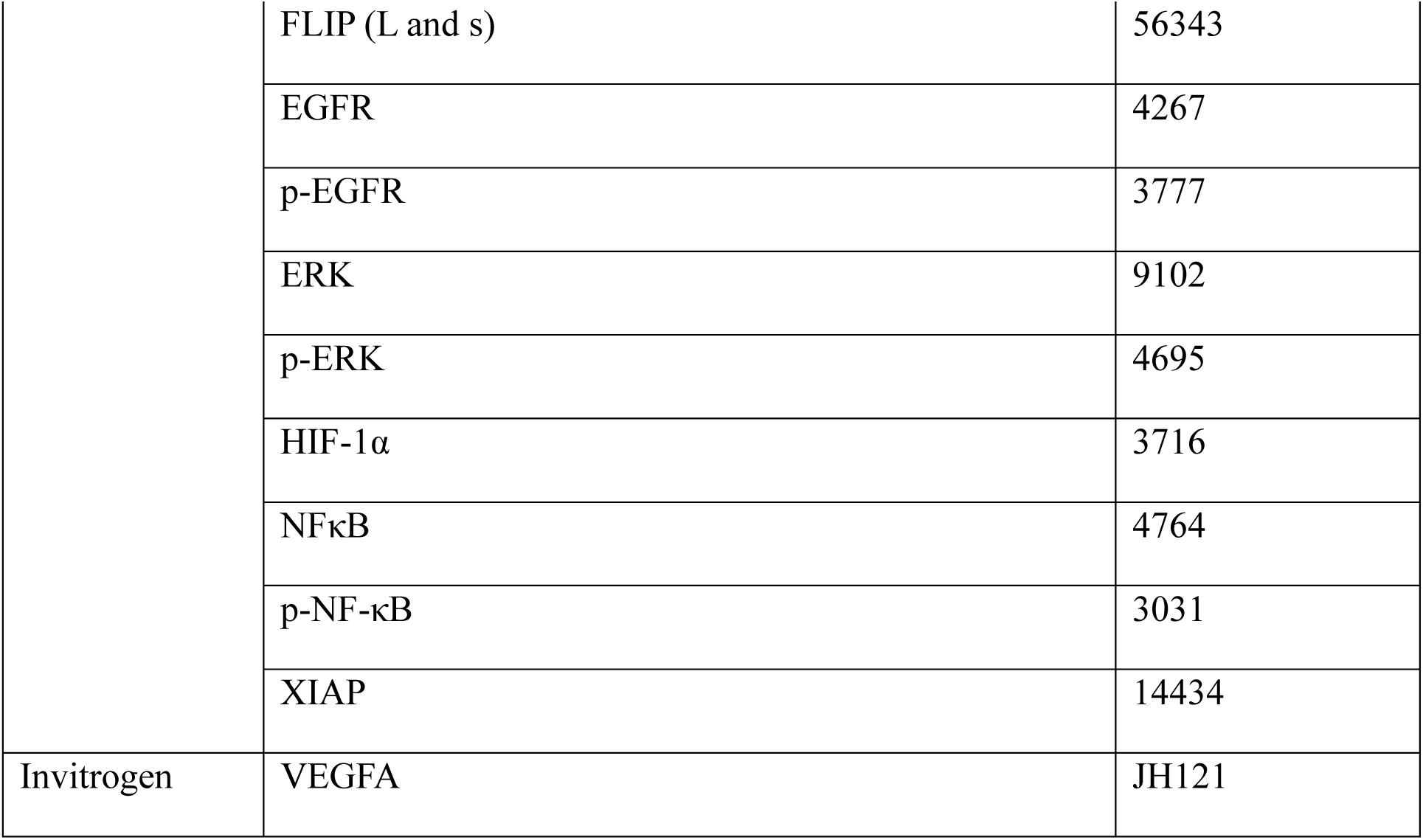
Vendor Information for qPCR and Antibodies Used in Western Blotting.

## REFERENCES

1 Barsouk, A., Aluru, J. S., Rawla, P., Saginala, K. & Barsouk, A. Epidemiology, Risk Factors, and Prevention of Head and Neck Squamous Cell Carcinoma. Med Sci (Basel) 11 (2023). 10.3390/medsci11020042

2 Tan, Y. et al. Oral squamous cell carcinomas: state of the field and emerging directions. Int J Oral Sci 15, 44 (2023). 10.1038/s41368-023-00249-w

3 Bose, P. et al. Tumor cell apoptosis mediated by cytoplasmic ING1 is associated with improved survival in oral squamous cell carcinoma patients. Oncotarget 5, 3210–3219 (2014). 10.18632/oncotarget.1907

4 Imbesi Bellantoni, M., et al. Oral Cavity Squamous Cell Carcinoma: An Update of the Pharmacological Treatment. Biomedicines 11 (2023). 10.3390/biomedicines11041112

5 Nokovitch, L. et al. Oral Cavity Squamous Cell Carcinoma Risk Factors: State of the Art. J Clin Med 12 (2023). 10.3390/jcm12093264

6 Wu, F., Du, Y., Hou, X. & Cheng, W. A prognostic model for oral squamous cell carcinoma using 7 genes related to tumor mutational burden. BMC Oral Health 22, 152 (2022). 10.1186/s12903-022-02193-3

7 D’Onofrio, I., Nardone, V., Reginelli, A. & Cappabianca, S. Chemoradiotherapy for Head and Neck Cancer. Cancers (Basel) 15 (2023). 10.3390/cancers15102820

8 De Felice, F., Cattaneo, C. G. & Franco, P. Radiotherapy and Systemic Therapies: Focus on Head and Neck Cancer. Cancers (Basel) 15 (2023). 10.3390/cancers15174232

9 Lingen, M. W. et al. Evidence-based clinical practice guideline for the evaluation of potentially malignant disorders in the oral cavity: A report of the American Dental Association. J Am Dent Assoc 148, 712–727 e710 (2017). 10.1016/j.adaj.2017.07.032

10 Micalizzi, D. S., Ebright, R. Y., Haber, D. A. & Maheswaran, S. Translational Regulation of Cancer Metastasis. Cancer Res 81, 517–524 (2021). 10.1158/0008-5472.CAN-20-2720

11 Song, P., Yang, F., Jin, H. & Wang, X. The regulation of protein translation and its implications for cancer. Signal Transduct Target Ther 6, 68 (2021). 10.1038/s41392-020-00444-9

12 Lee, L. J. et al. Cancer Plasticity: The Role of mRNA Translation. Trends Cancer 7, 134–145 (2021). 10.1016/j.trecan.2020.09.005

13 Deng, X., Yu, Y. V. & Jin, Y. N. Non-canonical translation in cancer: significance and therapeutic potential of non-canonical ORFs, m(6)A-modification, and circular RNAs. Cell Death Discov 10, 412 (2024). 10.1038/s41420-024-02185-y

14 Mahe, M., Rios-Fuller, T., Katsara, O. & Schneider, R. J. Non-canonical mRNA translation initiation in cell stress and cancer. NAR Cancer 6, zcae026 (2024). 10.1093/narcan/zcae026

15 Walters, B. & Thompson, S. R. Cap-Independent Translational Control of Carcinogenesis. Front Oncol 6, 128 (2016). 10.3389/fonc.2016.00128

16 Bressler, K. R. et al. Depletion of eukaryotic initiation factor 5B (eIF5B) reprograms the cellular transcriptome and leads to activation of endoplasmic reticulum (ER) stress and c-Jun N-terminal kinase (JNK). Cell Stress Chaperones 26, 253–264 (2021). 10.1007/s12192-020-01174-1

17 Pisareva, V. P. & Pisarev, A. V. eIF5 and eIF5B together stimulate 48S initiation complex formation during ribosomal scanning. Nucleic Acids Res 42, 12052–12069 (2014). 10.1093/nar/gku877

18 Harris, M. T. & Marr, M. T., 2nd. The intrinsically disordered region of eIF5B stimulates IRES usage and nucleates biological granule formation. Cell Rep 42, 113283 (2023). 10.1016/j.celrep.2023.113283

19 Ross, J. A. et al. Eukaryotic initiation factor 5B (eIF5B) provides a critical cell survival switch to glioblastoma cells via regulation of apoptosis. Cell Death Dis 10, 57 (2019). 10.1038/s41419-018-1283-5

20 Goldman, M. J. et al. Visualizing and interpreting cancer genomics data via the Xena platform. Nat Biotechnol 38, 675–678 (2020). 10.1038/s41587-020-0546-8

21 Huang, C. et al. Proteogenomic insights into the biology and treatment of HPV-negative head and neck squamous cell carcinoma. Cancer Cell 39, 361–379 e316 (2021). 10.1016/j.ccell.2020.12.007

22 Puram, S. V. et al. Single-Cell Transcriptomic Analysis of Primary and Metastatic Tumor Ecosystems in Head and Neck Cancer. Cell 171, 1611–1624 e1624 (2017). 10.1016/j.cell.2017.10.044

23 Satija, R., Farrell, J. A., Gennert, D., Schier, A. F. & Regev, A. Spatial reconstruction of single-cell gene expression data. Nat Biotechnol 33, 495–502 (2015). 10.1038/nbt.3192

24 McShane, L. M. et al. REporting recommendations for tumor MARKer prognostic studies (REMARK). Nat Clin Pract Urol 2, 416–422 (2005).

25 Arora, R. et al. NCBP2 and TFRC are novel prognostic biomarkers in oral squamous cell carcinoma. Cancer Gene Ther 30, 752–765 (2023). 10.1038/s41417-022-00578-8

26 De la Cruz-Morcillo, M. A. et al. p75 neurotrophin receptor and pro-BDNF promote cell survival and migration in clear cell renal cell carcinoma. Oncotarget 7, 34480–34497 (2016). 10.18632/oncotarget.8911

27 Ho, J. J. D. et al. Proteomics reveal cap-dependent translation inhibitors remodel the translation machinery and translatome. Cell Rep 37, 109806 (2021). 10.1016/j.celrep.2021.109806

28 DeCicco-Skinner, K. L. et al. Endothelial cell tube formation assay for the in vitro study of angiogenesis. J Vis Exp, e51312 (2014). 10.3791/51312

29 Panda, A. C., Martindale, J. L. & Gorospe, M. Polysome Fractionation to Analyze mRNA Distribution Profiles. Bio Protoc 7 (2017). 10.21769/BioProtoc.2126

30 Beilsten-Edmands, V. et al. eIF2 interactions with initiator tRNA and eIF2B are regulated by post-translational modifications and conformational dynamics. Cell Discov 1, 15020 (2015). 10.1038/celldisc.2015.20

31 Kim, E. et al. eIF2A, an initiator tRNA carrier refractory to eIF2alpha kinases, functions synergistically with eIF5B. Cell Mol Life Sci 75, 4287–4300 (2018). 10.1007/s00018-018-2870-4

32 Thakor, N. & Holcik, M. IRES-mediated translation of cellular messenger RNA operates in eIF2alpha- independent manner during stress. Nucleic Acids Res 40, 541–552 (2012). 10.1093/nar/gkr701

33 Ho, J. J. D. et al. Oxygen-Sensitive Remodeling of Central Carbon Metabolism by Archaic eIF5B. Cell Rep 22, 17–26 (2018). 10.1016/j.celrep.2017.12.031

34 Chukka, P. A. R., Wetmore, S. D. & Thakor, N. Established and Emerging Regulatory Roles of Eukaryotic Translation Initiation Factor 5B (eIF5B). Front Genet 12, 737433 (2021). 10.3389/fgene.2021.737433

35 Kearse, M. G. & Wilusz, J. E. Non-AUG translation: a new start for protein synthesis in eukaryotes. Genes Dev 31, 1717–1731 (2017). 10.1101/gad.305250.117

36 Lakshminarayana, S. et al. Molecular pathways of oral cancer that predict prognosis and survival: A systematic review. J Carcinog 17, 7 (2018). 10.4103/jcar.JCar_17_18

37 Johnson, D. E. et al. Head and neck squamous cell carcinoma. Nat Rev Dis Primers 6, 92 (2020). 10.1038/s41572-020-00224-3

38 Wang, Z. G. et al. eIF5B increases ASAP1 expression to promote HCC proliferation and invasion. Oncotarget 7, 62327–62339 (2016). 10.18632/oncotarget.11469

39 Magar, A. G., Morya, V. K., Kwak, M. K., Oh, J. U. & Noh, K. C. A Molecular Perspective on HIF-1alpha and Angiogenic Stimulator Networks and Their Role in Solid Tumors: An Update. Int J Mol Sci 25 (2024). 10.3390/ijms25063313

40 Palazon, A. et al. An HIF-1alpha/VEGF-A Axis in Cytotoxic T Cells Regulates Tumor Progression. Cancer Cell 32, 669–683 e665 (2017). 10.1016/j.ccell.2017.10.003

41 Hao, P. et al. Eukaryotic translation initiation factors as promising targets in cancer therapy. Cell Commun Signal 18, 175 (2020). 10.1186/s12964-020-00607-9

42 Ross, J. A. et al. Eukaryotic initiation factor 5B (eIF5B) regulates temozolomide-mediated apoptosis in brain tumour stem cells (BTSCs). Biochem Cell Biol 98, 647–652 (2020). 10.1139/bcb-2019-0329

43 Yuan, Y. et al. The translation initiation factor eIF3i up-regulates vascular endothelial growth factor A, accelerates cell proliferation, and promotes angiogenesis in embryonic development and tumorigenesis. J Biol Chem 289, 28310–28323 (2014). 10.1074/jbc.M114.571356

44 Young, R. M. et al. Hypoxia-mediated selective mRNA translation by an internal ribosome entry site-independent mechanism. J Biol Chem 283, 16309–16319 (2008). 10.1074/jbc.M710079200

